# Sex-Specific Evolution of the Genome-wide Recombination Rate

**DOI:** 10.1101/2020.07.08.194191

**Authors:** April L. Peterson, Bret A. Payseur

**Affiliations:** University of Wisconsin-Madison, Laboratory of Genetics

## Abstract

Although meiotic recombination is required for successful gametogenesis in most species that reproduce sexually, the rate of crossing over varies among individuals. Differences in recombination rate between females and males are perhaps the most striking form of this variation. To determine how sex shapes the evolution of recombination, we directly compared the genome-wide recombination rate in females and males across a common set of genetic backgrounds in house mouse. Our results reveal highly discordant evolutionary trajectories in the two sexes. Whereas male recombination rates show rapid evolution over short timescales, female recombination rates measured in the same strains are mostly static. Strains with high recombination in males have more double-strand breaks and stronger crossover interference than strains with low recombination in males, suggesting that these factors contribute to the sex-specific evolution we document. Our findings provide the strongest evidence yet that sex is a primary driver of recombination rate evolution.

## INTRODUCTION

Meiosis converts diploid germ cells into haploid gametes. During meiosis I, DNA crossovers aid the separation of homologous chromosomes by physically linking them and establishing tension between them on the spindle (Petronczki et al., 2003). The wrong number of recombination events can disrupt chromosomal segregation, leading to infertility, miscarriage, and birth defects (Hassold and Hunt, 2001). Recombination also shapes evolution by shuffling the combinations of genetic variants offspring inherit. Recombination affects the fates of beneficial and deleterious mutations (Felsenstein, 1974; Fisher, 1930; Hill and Robertson, 1966) and interacts with natural selection to leave gradients in genomic patterns of diversity (Begun and Aquadro, 1992; Charlesworth et al., 1993; Cutter and Payseur, 2013; Nachman and Payseur, 2012; Smith and Haigh, 1974).

The role of recombination in facilitating meiotic chromosome assortment suggests that the total number of crossovers in a cell – the genome-wide recombination rate – is an important cellular characteristic connected to organismal fitness. The dual pressures of ensuring at least one crossover per chromosome and minimizing levels of DNA damage and ectopic exchange are thought to impose lower and upper thresholds on the genome-wide recombination rate (Inoue and Lupski, 2002; Nagaoka et al., 2012). Yet, within these bounds, individuals from the same species can vary substantially in crossover number (Gruhn et al., 2013; Johnston et al., 2016; Kong et al., 2008; Ma et al., 2015).

Sex is perhaps the most notable axis along which recombination rate varies. Broadly speaking, sexual dimorphism in the genome-wide recombination rate assumes two forms. In species such as *Drosophila melanogaster*, one sex completes meiosis without forming crossovers (“achiasmy”), while the other sex recombines (Burt et al., 1991; Haldane, 1922; Huxley, 1928). Alternatively, in most species with recombination, crossovers occur in both sexes but at different rates (“heterochiasmy”). In these species, females tend to recombine more than males (Bell, 1982; Brandvain and Coop, 2012; Burt et al., 1991; Lenormand and Dutheil, 2005; Lorch, 2005). In plants, heterochiasmy is correlated with the opportunity for haploid selection (Lenormand and Dutheil, 2005).

Despite the establishment of these interspecific trends, an understanding of how sex shapes the evolution of recombination cannot be achieved with available data. Comprehensive comparisons of variation in female and male recombination rates within species have come from outbred populations of humans (Gruhn et al., 2013; Halldorsson et al., 2019; Kong et al., 2004, 2014, 2008), dog (Campbell et al., 2016), cattle (Ma et al., 2015; Shen et al., 2018), and Soay sheep (Johnston et al., 2016), in which the role of sex is confounded with the contributions of genetic variation. Although it is known that the level and direction of heterochiasmy can differ among species (Brandvain and Coop, 2012; Lenormand and Dutheil, 2005), the correlation between female and male recombination rates among closely related species remains poorly documented. Direct contrasts between the two sexes across a common, diverse set of genomic backgrounds that represent recent timescales would reveal whether the genome-wide recombination rate evolves differently in males and females.

Examining variation in the total number of crossovers in a sex-specific manner could also illuminate evolutionary connections between recombination rate and crossover positioning. Analyses of meiotic chromosome morphology in *Arabidopsis thaliana*, *Caenorhabditis elegans*, and *Mus musculus* suggest that the sex with more recombination usually has longer chromosome axes (Cahoon and Libuda, 2019). A survey of 51 species found conserved sex differences in the recombination landscape, including telomere-biased placement of crossovers in males but not in females (Sardell and Kirkpatrick, 2020). The degree to which a crossover reduces the probability of another crossover nearby (crossover interference) also differs between females and males (Otto and Payseur, 2019).

The house mouse, *Mus musculus*, is a compelling system for understanding how sex affects the evolution of recombination. Multiple subspecies share a most recent common ancestor approximately 0.5 million years ago (Geraldes et al., 2011), providing the opportunity to examine natural variation on recent evolutionary timescales. Wild *Mus musculus* belong to the same species as classical inbred strains of mice, where the molecular and cellular pathways that lead to crossovers have been studied extensively (Baudat et al., 2013; Bolcun-Filas and Schimenti, 2012; Handel and Schimenti, 2010). Single-cell immunofluorescent approaches make it possible to estimate genome-wide recombination rates in individual males and females (Koehler et al., 2002; Peters et al., 1997). A collection of wild-derived inbred strains founded from a variety of geographic locations is available, enabling genetic variation in recombination to be profiled across the species range. Most importantly, by measuring recombination rates in females and males from the same set of wild-derived inbred strains, the evolutionary dynamics of recombination can be directly compared in the two sexes.

In this paper, we report genome-wide recombination rates from both sexes in a diverse panel of wild-derived inbred strains of house mice and their close relatives. We demonstrate that recombination rate evolves differently in females and males, even over short timescales.

## RESULTS

### Genome-wide recombination rate evolves differently in females and males

We used counts of MLH1 foci per cell to estimate genome-wide recombination rates in 14 wild-derived inbred strains sampled from three subspecies of house mice (*M. musculus domesticus*, *M. m. musculus* and *M. m. molossinus*) and three other species of Mus (*M. spretus*, *M. spicilegus*, and *M. caroli*). Mean MLH1 focus counts for 188 mice were quantified from an average of 21.77 spermatocytes per male (for a total of 1,742 spermatocytes) and 17.85 oocytes per female (for a total of 1,427 oocytes) (Table 1).

**Table 1.**
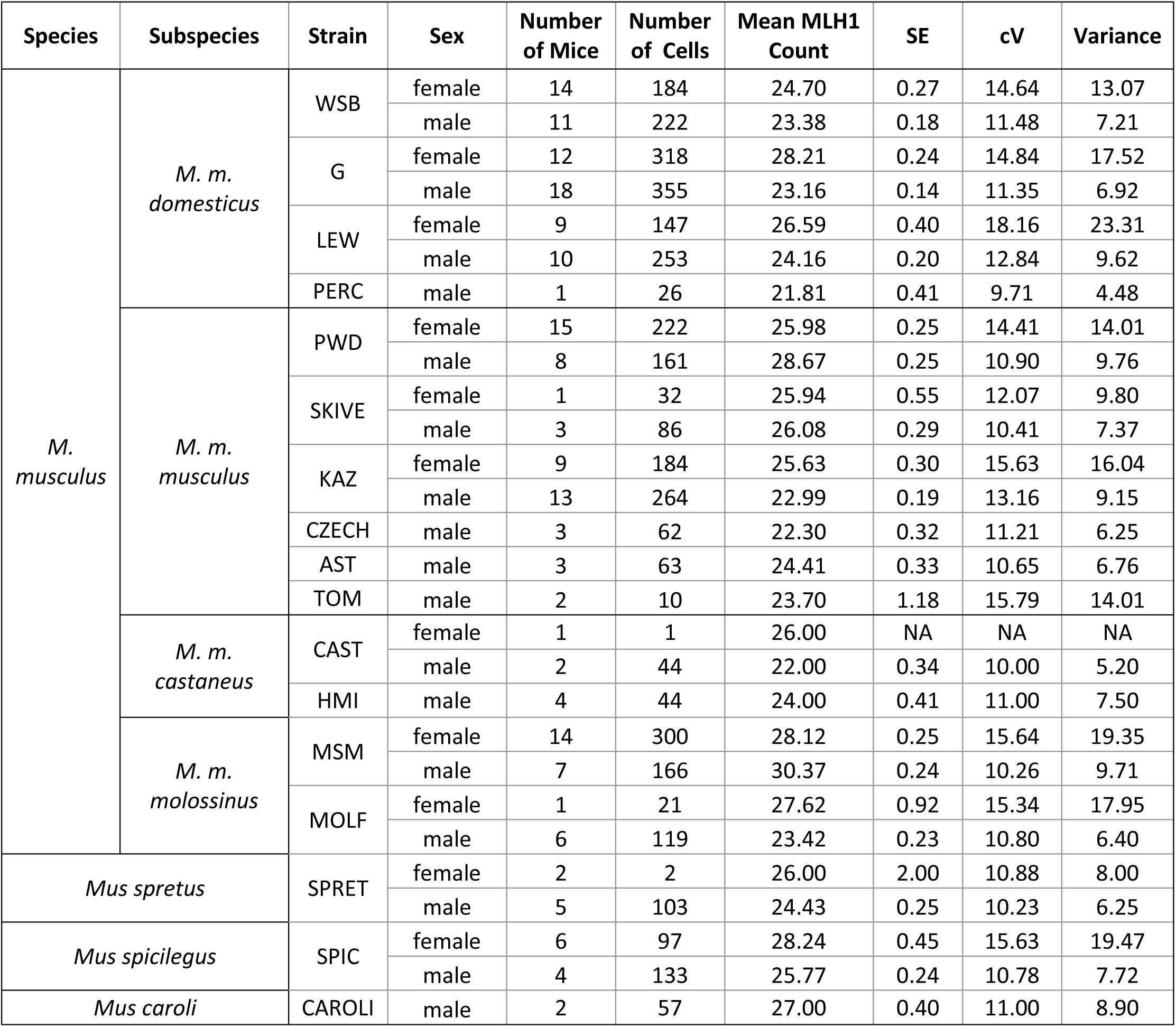

| Species | Subspecies | Strain | Sex | Number of Mice | Number of Cells | Mean MLH1 Count | SE | cV | Variance |
| --- | --- | --- | --- | --- | --- | --- | --- | --- | --- |
| M. musculus | M. m. domesticus | WSB | female | 14 | 184 | 24.70 | 0.27 | 14.64 | 13.07 |
|  |  |  | male | 11 | 222 | 23.38 | 0.18 | 11.48 | 7.21 |
|  |  | G | female | 12 | 318 | 28.21 | 0.24 | 14.84 | 17.52 |
|  |  |  | male | 18 | 355 | 23.16 | 0.14 | 11.35 | 6.92 |
|  |  | LEW | female | 9 | 147 | 26.59 | 0.40 | 18.16 | 23.31 |
|  |  |  | male | 10 | 253 | 24.16 | 0.20 | 12.84 | 9.62 |
|  |  | PERC | male | 1 | 26 | 21.81 | 0.41 | 9.71 | 4.48 |
|  | M. m. musculus | PWD | female | 15 | 222 | 25.98 | 0.25 | 14.41 | 14.01 |
|  |  |  | male | 8 | 161 | 28.67 | 0.25 | 10.90 | 9.76 |
|  |  | SKIVE | female | 1 | 32 | 25.94 | 0.55 | 12.07 | 9.80 |
|  |  |  | male | 3 | 86 | 26.08 | 0.29 | 10.41 | 7.37 |
|  |  | KAZ | female | 9 | 184 | 25.63 | 0.30 | 15.63 | 16.04 |
|  |  |  | male | 13 | 264 | 22.99 | 0.19 | 13.16 | 9.15 |
|  |  | CZECH | male | 3 | 62 | 22.30 | 0.32 | 11.21 | 6.25 |
|  |  | AST | male | 3 | 63 | 24.41 | 0.33 | 10.65 | 6.76 |
|  |  | TOM | male | 2 | 10 | 23.70 | 1.18 | 15.79 | 14.01 |
|  | M. m. castaneus | CAST | female | 1 | 1 | 26.00 | NA | NA | NA |
|  |  |  | male | 2 | 44 | 22.00 | 0.34 | 10.00 | 5.20 |
|  |  | HMI | male | 4 | 44 | 24.00 | 0.41 | 11.00 | 7.50 |
|  | M. m. molossinus | MSM | female | 14 | 300 | 28.12 | 0.25 | 15.64 | 19.35 |
|  |  |  | male | 7 | 166 | 30.37 | 0.24 | 10.26 | 9.71 |
|  |  | MOLF | female | 1 | 21 | 27.62 | 0.92 | 15.34 | 17.95 |
|  |  |  | male | 6 | 119 | 23.42 | 0.23 | 10.80 | 6.40 |
| Mus spretus |  | SPRET | female | 2 | 2 | 26.00 | 2.00 | 10.88 | 8.00 |
|  |  |  | male | 5 | 103 | 24.43 | 0.25 | 10.23 | 6.25 |
| Mus spicilegus |  | SPIC | female | 6 | 97 | 28.24 | 0.45 | 15.63 | 19.47 |
|  |  |  | male | 4 | 133 | 25.77 | 0.24 | 10.78 | 7.72 |
| Mus caroli |  | CAROLI | male | 2 | 57 | 27.00 | 0.40 | 11.00 | 8.90 |

Graphical comparisons reveal sex-specific dynamics to the evolution of genome-wide recombination rate (Figure 1A). First, MLH1 focus counts differ between females and males in most strains. Second, the difference in counts between the sexes varies among strains. Although most strains show more MLH1 foci in females, two strains (*musculus^PWD^* and *molossinus^MSM^*) exhibit higher counts in males. In females, numbers of MLH1 foci are evenly distributed around the sex-wide mean of approximately 25 (Figure 1B). In stark contrast, males largely separate into two groups of strains with high numbers (near 30) and low numbers (near 23) of foci (Figure 1C). Strain mean MLH1 focus counts from females and males are uncorrelated (Spearman’s ρ = 0.08; p = 0.84) across the set of strains.

**Figure 1.**
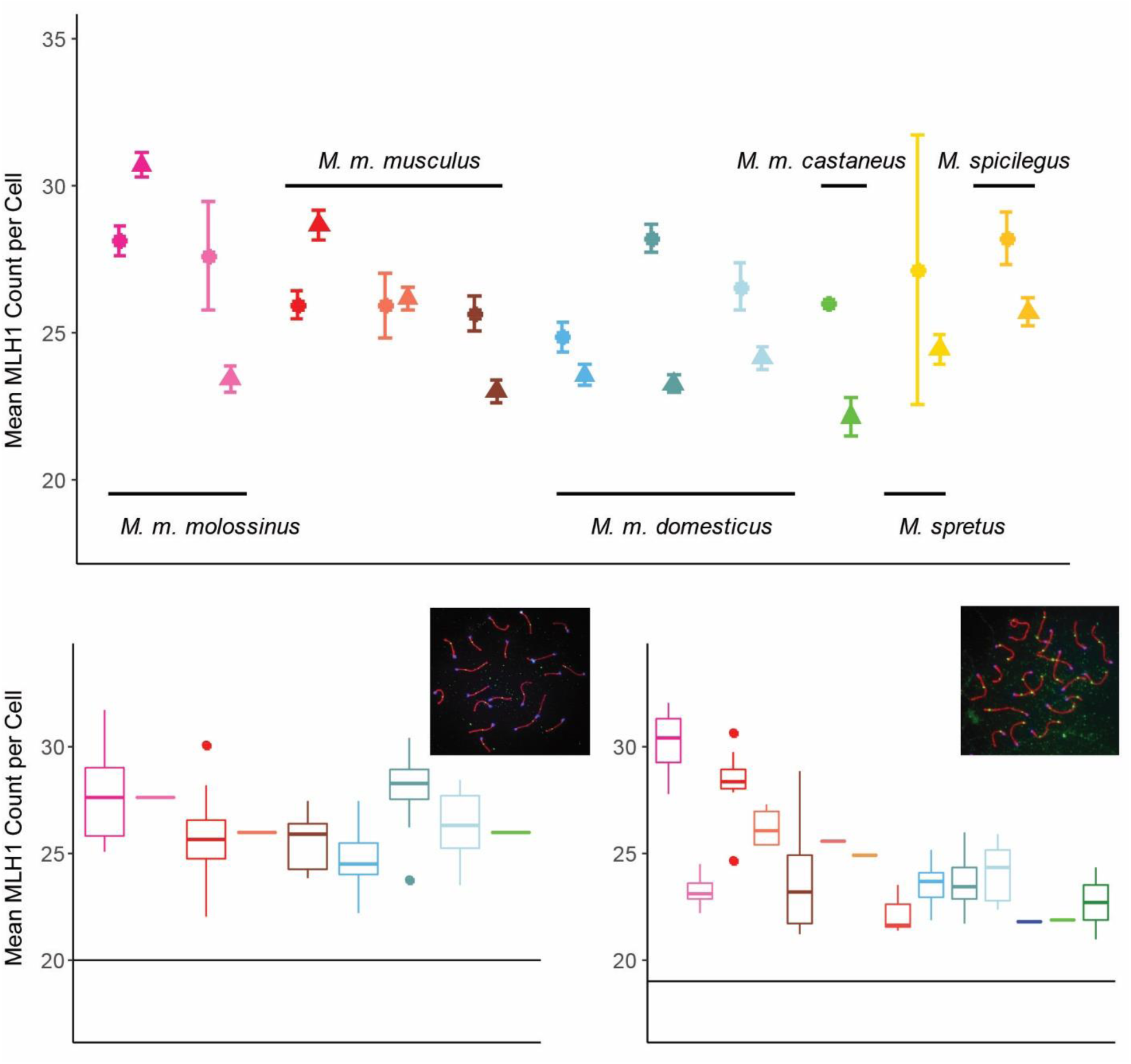
MLH1 Counts. A) Strain mean MLH1 counts (+/- 2 standard errors) in both sexes. Females = circles; males = triangles. B) Boxplots of female MLH1 counts for strains of house mice. Whiskers indicate interquartile range. Inset: example oocyte, SYCP3 stained in red, CREST (centromeres) stained in blue and MLH1 foci stained in green. Horizontal line at 20 indicates the expected minimum number of foci per cell. C) Boxplots of male MLH1 counts for strains of house mice. Inset: example spermatocyte. Additional strains with only male observations are included with the values from Table 2.

**Table 2.**

| Model | Dataset(s) | Dependent Variable(s) | Fixed Effects | Random Effects |
| --- | --- | --- | --- | --- |
| M1 | females and males from 8 strains | mouse average | Subspecies<br>Sex<br>Subspecies*Sex | Strain |
| M2 | females and males from 8 strains | mouse average | Subspecies<br>Sex<br>Strain<br>Subspecies*Sex<br>Subspecies*Strain<br>Sex*Strain |  |
| M3 | females and males from 8 strains | mouse average | Sex<br>Strain<br>Sex*Strain |  |
| M4 | females from 8 strains | female mouse average | Subspecies<br>Strain<br>Subspecies*Strain |  |
| M4 | males from 12 strains | male mouse average | Subspecies<br>Strain<br>Subspecies*Strain |  |
| M5 | females from 8 strains | female mouse average | Strain |  |
| M5 | males from 12 strains | male mouse average | Strain |  |

To further partition variation in recombination rate, we fit a series of linear models to mean MLH1 focus counts from 137 house mice from *M. m. domesticus*, *M. m. musculus* and *M. m. molossinus* (Table 2; detailed results available in Supplemental Tables 1-7). Strain, sex, subspecies, and sex*subspecies each affect MLH1 focus count in a linear mixed model (M1; strain (random effect): p < 10^-4^; sex: p = 3.64 x 10^-6^; subspecies: p = 9.69 x 10^-4^; subspecies*sex: p = 1.8 x 10^-4^).

The effect of subspecies is no longer significant in a model treating all factors as fixed effects (M2; *musculus* p = 0.24, *molossinus* p = 0.1), highlighting strain and sex as salient variables. Two strains exhibit strong effects on MLH1 focus count (M3; *domesticus^G^* p = 1.78 x 10^-6^; *domesticus^LEW^* p = 0.02), with sex-strain interactions involving three strains (M3; *domesticus^G^* p < 10^-6^; *molossinus^MSM^* p < 10^-6^; *musculus^PWD^* p = 3.87 x 10^-4^).

In separate analyses of males (M4; n = 71), three strains disproportionately shape MLH1 focus count (as observed in Figure 1C): *musculus^PWD^* (p = 3.6 x 10^-7^; effect = 6.11 foci, *molossinus^MSM^* (p = 6.3 x 10^-9^; effect = 6.91), and *musculus^SKIVE^* (p = 8.22 x 10^-4^; effect = 4.04). These three strains point to substantial evolution in the genome-wide recombination rate in spermatocytes; we subsequently refer to them as “high-recombination” strains. In females (M4; n= 76), three strains affect MLH1 focus count: *domesticus^G^* (p = 8.7 x 10^-6^; effect = 3.3), *molossinus^MSM^* (p = 2.43 x 10^-5^; effect = 2.99), and *domesticus^LEW^* (p = 0.03; effect = 1.69). Strain effect sizes in females are modest in magnitude compared to those in males. Together, these results demonstrate that the genome-wide recombination rate evolves in a highly sex-specific manner.

### Synaptonemal complexes are longer in females

The variation in sex differences in recombination we discovered provided an opportunity to determine whether sex differences in chromatin compaction, as measured by the length of the synaptonemal complex (SC), are reversed when heterochiasmy is reversed. In all strains except *musculus^SKIVE^*, females have longer SCs than males, whether SC length was estimated as the total length across bivalents or as the length of short bivalents (t-tests; all p < 0.05, except short bivalents in *musculus^SKIVE^*, p = 0.11). Among short bivalents (to which the female X bivalent does not contribute), female to male ratios of mouse mean SC length range from 1.26 (*musculus^PWD^*) to 1.52 (*domesticus^WSB^*) across strains. That females have longer SCs is further supported by models that include covariates, which identify sex as the most consistently significant effect for total SC length (M1: p = 2.56 x 10^-31^; M2: p = 2.56 x 10^-8^; M3: p = 2.56 x 10^-8^) (Supplemental Tables 8-14) and short bivalent SC length (M1: p = 1.12 x 10^-11^; M2: p < 10^-6^; M3: p < 1.33 x 10^-7^) (Supplemental Tables 15-21). The existence of some subspecies and strain effects on total SC length and short bivalent SC length further indicates that SC length has evolved among strains and among subspecies.

In summary, two approaches for measuring SC length demonstrate that females have longer SCs (chromosome axes), even in strains in which males recombine more. This pattern implies that in high-recombination strains, spermatocytes have less space than oocytes in which to position additional crossovers.

### Females and males differ in crossover positions and crossover interference

We used normalized positions of MLH1 foci along bivalents with a single focus to compare crossover location while controlling for differences in SC length. In all strains, MLH1 foci tend to be closer to the telomere in males (mean normalized position in males: 0.68; mean normalized position in females: 0.56; paired t-test; p = 8.49 x 10^-4^). Sex is also the strongest determinant of MLH1 focus position in the models we tested (M1: p = 2.82 x 10^-26^; M2: p = 3.96 x 10^-8^; M3: p = 3.96 x 10^-8^) (Supplemental Tables 24-30).

Males have longer normalized mean inter-focal distances (IFD_norm_) than females in seven out of eight strains (t-tests; p < 0.05), with only *musculus^KAZ^* showing no difference (p = 0.33). Examination of IFD_norm_ distributions indicates that females are centered at approximately 50% and show a slight enrichment of low (<25%) values, whereas males are enriched for higher values. Models treating IFD_norm_ as the dependent variable support the inference of stronger interference in males, with sex being the most significant variable (M1: p = 9.08 x 10^-12^; M2: p = 0.01; M3: p = 0.01) (Supplemental Tables 34-37). In contrast, there is no clear signal of sex differences in raw mean inter-focal distances (IFD_raw_) (Supplemental Tables 38-40) across the full set of strains, whether they are considered separately or together. Visualization of normalized MLH1 foci positions on bivalents with two crossovers (Figure 3; Supplemental Figure 3) further suggests that interference distances vary more in females than in males, and that males display a stronger telomeric bias in the placement of the distal crossover.

**Figure 2.**
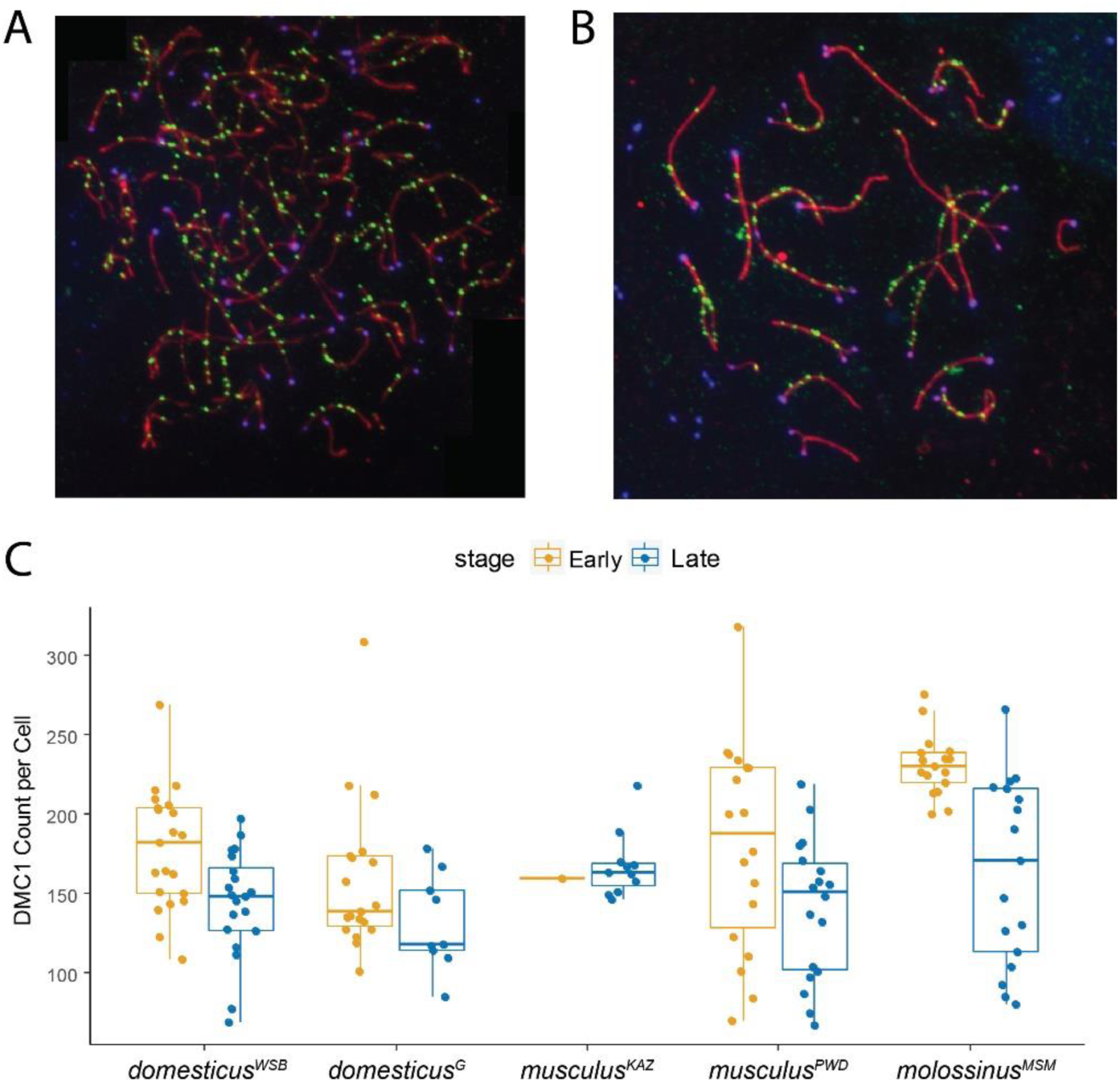
DMC1 Counts in Males. A) Example early zygotene spermatocyte spread. SYCP3 stained in red, CREST (centromeres) stained in blue and DMC1 stained in green. B) Example late zygotene spermatocyte spread. C) Boxplots of DMC1 counts for strains of house mice. Whiskers indicate interquartile range.

**Figure 3.**
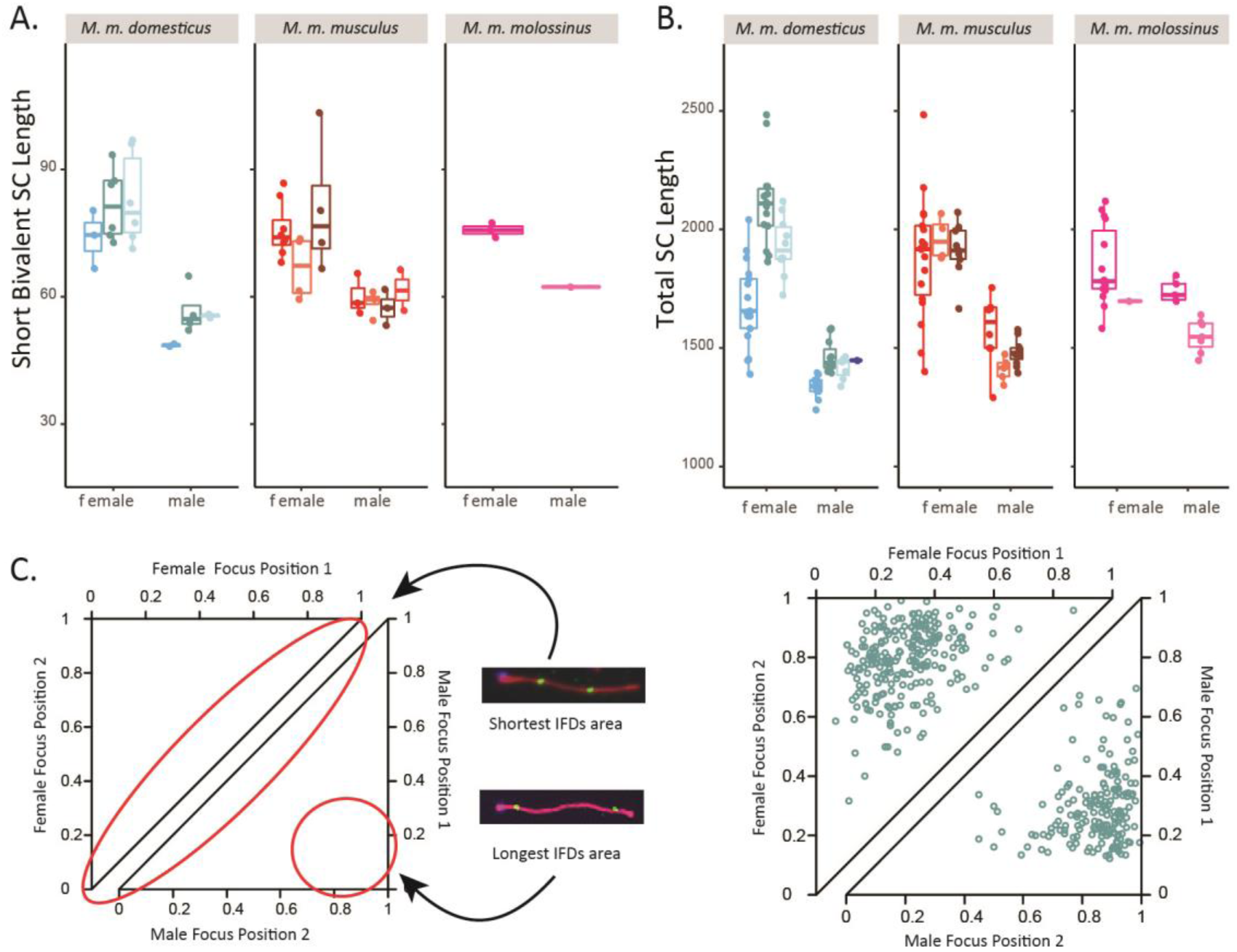
Sex Differences in Synaptonemal Complex (SC) Length and MLH1 Foci Positions. A) Mouse average SC length of short bivalents. Whiskers indicate interquartile range. B) Mouse average total SC length. C) Example of sex differences in inter-focal distances and foci locations on bivalents with two foci. Female observations shown in top triangle; male observations shown in bottom triangle. Data from domesticus^G^.

In summary, controlling for differences in SC length (chromatin compaction) indicates that interference is consistently stronger in males, whereas interference on the physical scale is similar in the two sexes.

### Evolution of genome-wide recombination rate is dispersed across bivalents, associated with double-strand break number, and connected to crossover interference

We used the contrast between males from high-recombination strains and males from low-recombination strains to identify features of the recombination landscape associated with evolutionary transitions in the genome-wide recombination rate. We considered proportions of bivalents with different numbers of crossovers, double-strand break number, SC length, and crossover positioning.

Ninety-six percent of single bivalents in our pooled dataset (n = 9,569) have either one or two MLH foci (Supplemental Figure 2). The proportions of single-focus (1CO) bivalents vs. double-focus (2CO) bivalents distinguish high-recombination strains from low-recombination strains (Supplemental Figure 2). High-recombination strains are enriched for 2CO bivalents at the expense of 1CO bivalents: proportions of 2CO bivalents are 0.33 in *musculus^SKIVE^*, 0.44 in *musculus^PWD^*, and 0.51 in *molossinus^MSM^* (Supplemental Figure 3). Following patterns in the genome-wide recombination rate, male *musculus^PWD^* and male *molossinus^MSM^* have 2CO proportions that are more similar to each other than to strains from their own subspecies (chi-square tests; *musculus^PWD^* vs. *molossinus^MSM^*: p = 0.37; *musculus^PWD^* vs. *musculus^KAZ^*: p = 1.23 x 10^-31^; *molossinus^MSM^* vs. *molossinus^MOLF^*: p = 2.34 x 10^-6^). These results demonstrate that evolution of the genome-wide recombination rate reflects changes in crossover number across multiple bivalents.

To begin to localize evolution of genome-wide recombination rate to steps of the recombination pathway, we counted DMC1 foci in prophase spermatocytes as markers for double-strand breaks (DSBs). DMC1 foci were counted in a total of 76 early zygotene and 75 late zygotene spermatocytes from two high-recombination strains (*musculus^PWD^* and *molossinus^MSM^*) and three low-recombination strains (*musculus^KAZ^*, *domesticus^WSB^*, and *domesticus^G^*) (Table 3). High-recombination strains have significantly more DMC1 foci than low-recombination strains in early zygotene cells (t-test; p < 10^-6^). In contrast, the two strain groups do not differ in DMC1 foci in late zygotene cells (t-test; p = 0.66). Since DSBs are repaired as either COs or non-crossovers (NCOs), the ratio of MLH1 foci to DMC1 foci can be used to estimate the proportion of DSBs designated as COs. High-recombination and low-recombination strains do not differ in the MLH1/DMC1 ratio, whether DMC1 foci were counted in early zygotene cells or late zygotene cells (t-test; p > 0.05). These results raise the possibility that the evolution of genome-wide recombination rate is primarily determined by processes that precede the CO/NCO decision, at least in house mice.

**Table 3.**

| Recombination Group | Strain | Early Zygotene |  |  | Late Zygotene |  |  |
| --- | --- | --- | --- | --- | --- | --- | --- |
|  |  | Cells | Mean | MLH1:DMC1 Ratio | Cells | Mean | MLH1:DMC1 Ratio |
| Low | *domesticus*<sup>WSB</sup> | 21 | 177.76 | 0.14 | 20 | 144.25 | 0.17 |
|  | *domesticus*<sup>G</sup> | 19 | 158.16 | 0.15 | 9 | 131.78 | 0.18 |
|  | *musculus*<sup>KAZ</sup> | 1 | 159.00 | 0.15 | 11 | 167.36 | 0.14 |
| High | *musculus*<sup>PWD</sup> | 18 | 180.22 | 0.16 | 18 | 140.78 | 0.21 |
|  | *molossinus*<sup>MSM</sup> | 17 | 231.00 | 0.14 | 17 | 164.41 | 0.19 |

Total SC length only partially differentiates high-recombination strains from low-recombination strains (Figure 3). Whereas high-recombination strains as a group have significantly greater total SC length than low-recombination strains (t-test; p = 0.01), separate tests within subspecies show that the two strain categories differ within *M. m. molossinus* (p = 2.59 x 10^-4^) but not within *M. m. musculus* (p = 0.65). Additionally, mouse means for the reduced (short and long) bivalent datasets do not differ between high-recombination and low-recombination strains (t-test; short: p = 0.84; long: p = 0.19). In a model with total SC length as the dependent variable (M4), the two subspecies effects are significant (*M. m. musculus* p = 3.95 x 10^-7^; *M. m. molossinus* p = 3.33 x 10^-7^), but there are also strain-specific effects (Supplemental Table 13). In models with SC lengths of short and long bivalents as dependent variables, several subspecies and strain effects reach significance (p < 0.05) (Supplemental Table 20,21, 22, and 23), but they are not consistent across models. Collectively, these results reveal that evolution of SC length is not strongly associated with evolution of genome-wide recombination rate in house mice.

In summary, evolution of the genome-wide recombination rate in males is connected to double-strand break number and crossover interference, but not to SC length and crossover position (on single-crossover bivalents).

## DISCUSSION

By comparing recombination rates in females and males from the same diverse set of genetic backgrounds, we isolated sex as a primary factor in the evolution of this fundamental meiotic trait. Recombination rate differences are more pronounced in males than females. Because inter-strain divergence times are identical for the two sexes, this observation demonstrates that the genome-wide recombination rate evolves faster in males. More generally, recombination rate divergence is decoupled in females and males. These disparities are remarkable given that recombination rates for the two sexes were measured in identical genomic backgrounds (other than the number and identity of sex chromosomes). Our results provide the strongest evidence yet that the genome-wide recombination rate follows distinct evolutionary trajectories in males and females.

At the genetic level, the sex-specific patterns we documented indicate that some mutations responsible for the evolution of recombination rate have dissimilar phenotypic effects in the two sexes. A subset of the genetic variants associated with genome-wide recombination rate within populations of humans (Kong et al., 2004, 2008, 2014; Halldorsson et al., 2019), Soay sheep (Johnston et al., 2016), and cattle (Ma et al., 2015; Shen et al., 2018) appear to show sex-specific properties, including opposite effects in females and males. Furthermore, inter-sexual correlations for recombination rate are weak in humans (Fledel-Alon et al., 2011) and Soay sheep (Johnston et al., 2016). Crosses between the strains we surveyed could be used to identify and characterize the genetic variants responsible for recombination rate evolution in house mice (Dumont and Payseur, 2011; Wang et al., 2019; Wang and Payseur, 2017). These variants could differentially affect females and males at any step in the recombination pathway. Although our DMC1 profiling was limited to males from a small number of strains (for practical reasons), our findings suggest that mutations that determine the number of double-strand breaks contribute to sex-specific evolution in the recombination rate. A study of two classical inbred strains and one wild-derived inbred strain of house mice also found a positive association between crossover number and double-strand break number in males (Baier et al., 2014).

Another implication of our results is that the connection between recombination rate and fitness differs between males and females. Little is known about whether and how natural selection shapes recombination rate in nature (Dapper and Payseur, 2017; Ritz et al., 2017). Samuk et al. (2020) recently used a quantitative genetic test to conclude that an 8% difference in genome-wide recombination rate between females from two populations of *Drosophila pseudoobscura* was caused by natural selection. Applying similar strategies to species in which both sexes recombine, including house mice, would be a logical next step to understanding the sex-specific evolution of recombination rate.

Population genetic models have been built to explain sexual dimorphism in the number and placement of crossovers, which is a common phenomenon (Brandvain and Coop, 2012; Sardell and Kirkpatrick, 2020). Modifier models predicted that lower recombination rates in males will result from haploid selection (Lenormand, 2003) or sexually antagonistic selection on coding and cis-regulatory regions of genes (Sardell and Kirkpatrick, 2020). Another modifier model showed that meiotic drive could stimulate female-specific evolution of the recombination rate (Brandvain and Coop, 2012). Although these models fit the conserved pattern of sex differences in crossover positions, they do not readily explain our observations of sex-specific evolution in the genome-wide recombination rate. In particular, the alternation across strains in which sex has more crossovers is unexpected.

We propose an alternative interpretation of our findings based on the cell biology of gametogenesis. During meiosis, achieving a stable chromosome structure requires the attachment of kinetochores to opposite poles of the cell and at least one crossover to create tension across the sister chromosome cohesion distal to chiasmata (Dumont and Desai, 2012; Lane and Kauppi, 2019; Subramanian and Hochwagen, 2014; VanVeen and Hawley, 2003). The spindle assembly checkpoint (SAC) prevents aneuploidy by ensuring that all bivalents are correctly attached to the microtubule spindle (“bi-oriented”) before starting the metaphase-to-anaphase transition via the release of the sister cohesion holding homologs together (Lane and Kauppi, 2019). Hence, selection seems likely to favor mutations that optimize the process of bi-orientation and chromosome separation, thereby prohibiting the SAC from delaying the cell cycle or triggering apoptosis. Multiple lines of evidence indicate that the SAC is more effective in spermatogenesis than in oogenesis (Lane and Kauppi, 2019), perhaps due to the presence of the acentrosome spindle (So et al., 2019) and larger cell volume (Kyogoku and Kitajima, 2017) in oocytes. The higher stringency of the SAC during spermatogenesis suggests that selection will be better at removing mutations that interfere with bi-orientation in males than in females. Therefore, faster male evolution of the genome-wide recombination rate could be driven by the more stringent SAC acting on chromosome structures at the metaphase I alignment.

Our SAC model is consistent with other features of our data. We showed that widespread sex differences in broad-scale crossover positioning (Sardell and Kirkpatrick, 2020) apply across house mice, even in lineages where the direction of heterochiasmy is reversed. Faster spermatogenesis may select for synchronization of the separation across all homologs within the cell (Kudo et al., 2009), whereas in oogenesis, the slower cell cycle and multiple arrest stages may require chromosome structures with greater stability on the meiosis I spindle, especially for those organisms that undergo dictyate arrest (Lee, 2019).

We propose that the SAC model also can explain the correlated evolution of stronger crossover interference and higher genome-wide recombination rate in male house mice. Our results show that crossovers are spaced further apart in strains enriched for double-crossover bivalents when SC length is considered and bivalent size effects are minimized. Assuming chromatin compaction between (prophase) pachytene and metaphase is uniform along bivalents, this increased spacing is expected to expand the area for sister cohesion to connect homologs and may improve the fidelity of chromosomal segregation. While the SAC model postulates direct fitness effects of interference, a modifier model predicted that indirect selection on recombination rate – via its modulation of offspring genotypes – can strengthen interference as well (Goldstein et al., 1993).

Regardless of the underlying mechanism, our results provide a rare demonstration that crossover interference can diverge over short evolutionary timescales. The notion that stronger interference can co-evolve with higher genome-wide recombination rate is supported by differences between breeds of cattle (Ma et al., 2015). In contrast, mammalian species with stronger interference tend to exhibit lower genome-wide recombination rates (Otto and Payseur, 2019; Segura et al., 2013). The evolution of crossover interference and its relationship to changes in crossover number on the genomic scale is a topic deserving of more empirical and theoretical work.

Our findings further reveal that evolution of the genome-wide recombination rate does not require major changes in the degree of chromatin compaction. Female house mice consistently show longer SCs, even in strains with more recombination in males. Studies in mice (Lynn et al., 2002; Petkov et al., 2007) and humans (Gruhn et al., 2013; Tease and Hulten, 2004) suggest that chromosomal axes are longer (and DNA loops are shorter) in females than males. Some authors have suggested that conserved sex differences in crossover positioning (more uniform placement in females) and interference strength (stronger interference in males) could be due to looser chromatin packing of the meiotic chromosome structure in females (Haenel et al., 2018; Petkov et al., 2007). A cellular model designed to explain interference attributes sexual dimorphism in chromatin structure to greater cell volumes and oscillatory movements of telomeres and kinetochores in oocytes (Hultén, 2011). Recent work in mice connects the sparser recombination landscape in females to sex differences in crossover maturation efficiency (Wang et al., 2017).

Our conclusions are accompanied by several caveats. First, MLH1 foci only identify interfering crossovers (Holloway et al., 2008). Although most crossovers belong to this class (Holloway et al., 2008), our approach likely underestimated genome-wide recombination rates. Evolution of the number of non-interfering crossovers is a subject worth examining. A second limitation is that our investigation of crossover locations was confined to the relatively low resolution possible with immunofluorescent cytology. Positioning crossovers with higher resolution could reveal additional evolutionary patterns. Finally, the panel of inbred lines we surveyed may not be representative of recombination rate variation within and between subspecies of house mice. We considered most available wild-derived inbred lines, but house mice have a broad geographic distribution. Nevertheless, we expect our primary conclusion that recombination rate evolves in a sex-specific manner to be robust to geographic sampling because differences between females and males exist for the same set of inbred strains.

While the causes of sex differences in recombination remain mysterious (Lenormand et al., 2016), our conclusions have implications for a wide range of recombination research. For biologists uncovering the cellular and molecular determinants of recombination, our results suggest that mechanistic differences between the sexes could vary by genetic background. For researchers charting the evolutionary trajectory of recombination, our findings indicate that sex-specific comparisons are crucial. For theoreticians building evolutionary models of recombination, different fitness regimes and genetic architectures in females and males should be considered. Elevating sex as a primary determinant of recombination would be a promising step toward integrating knowledge of cellular mechanisms with evolutionary patterns to understand recombination rate variation in nature.

## MATERIALS AND METHODS

### Mice

We used a panel of wild-derived inbred strains of house mice (*Mus musculus*) and related murid species to profile natural genetic variation in recombination (Table 4). Mice from the same inbred strain served as biological replicates. Our survey included 5 strains from *M. m. musculus*, 4 strains from *M. m. domesticus*, 2 strains from *M. m. molossinus*, 2 strains from *M. m. castaneus*, and 1 strain each from *M. spicilegus*, *M. spretus* and *M. caroli*. We subsequently denote strains by their abbreviated subspecies and name (*e.g. domesticus^WSB^*).

**Table 4.**
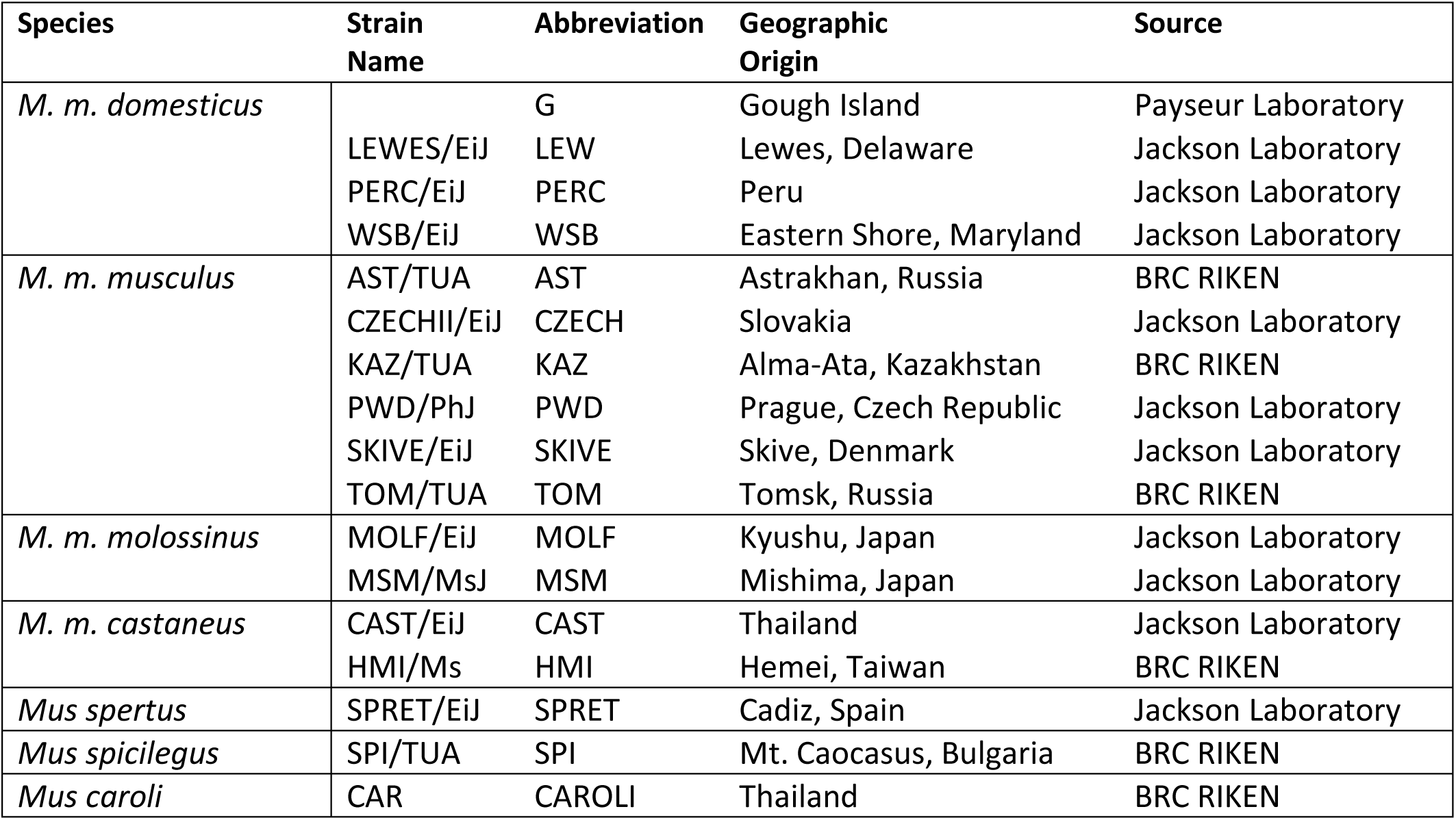

Mice were housed at dedicated, temperature-controlled facilities in the UW-Madison School of Medicine and Public Health, with the exception of mice from Gough Island, which were housed in a temperature-controlled facility in the UW-Madison School of Veterinary Medicine. Mice were sampled from a partially inbred strain of Gough Island mice, after approximately 6 generations of brother-sister matting. All mice were provided with *ad libitum* food and water. Procedures followed protocols approved by IACUC.

### Tissue Collection and Immunohistochemistry

The same dry-down spread technique was applied to both spermatocytes and oocytes, following Peters et al. (1997), with adjustment for volumes. Spermatocyte spreads were collected and prepared as described in Peterson et al. (2019). The majority of mice used for MLH1 counts were between 5 and 12 weeks of age. Juvenile males between 12 and 15 days of age were used for DMC1 counts. Both ovaries were collected from embryos (16-21 embryonic days) or neonates (0-48 hours after birth). Whole testes were incubated in 3ml of hypotonic solution for 45 minutes. Decapsulated ovaries were incubated in 300ul of hypotonic solution for 45 minutes. Fifteen microliters of cell slurry (masticated gonads) were transferred to 80ul of 2% PFA solution. Cells were fixed in this solution and dried in a humid chamber at room temperature overnight. The following morning, slides were treated with a Photoflow wash (Kodak, Rochester, NY, diluted 1:200). Slides were stored at −20*C if not stained immediately. To visualize the structure of meiotic chromosomes, we used antibody markers for the centromere (CREST) and lateral element of the synaptonemal complex (SC) (SYCP3). Crossovers (COs) were visualized as MLH1 foci. Double strand breaks (DSBs) were visualized as DMC1 foci. The staining protocol followed (Anderson et al., 1999) and (Koehler et al., 2002). Antibody staining and slide blocking were performed in 1X antibody dilution buffer (ADB) (normal donkey serum (Jackson ImmunoResearch, West Grove, PA), 1X PBS, bovine serum albumin (Sigma-Aldrich, St. Louis, MO), and Triton X-100 (Sigma-Aldrich, Stt. Louis, MO)). Following a 30-minute blocking wash in ABD, each slide was incubated with 60ul of a primary antibody master mix for 48 hours at 37*C. The master mix recipe contained polyclonal anti-rabbit anti-MLH1 (Calbiochem, San Diego CA; diluted 1:50) or anti-rabbit anti-DMC1 (mix of DMC1), anti-goat polyclonal anti-SYCP3, (Abcam, Cambridge, UK; diluted 1:50), and anti-human polyclonal antibody to CREST (Antibodies, Inc, Davies, CA; diluted 1:200) suspended in ADB. Slides were washed twice in 50ml ADB before the first round of secondary antibody incubation for 12 hours at 37*C. Alexa Fluor 488 donkey anti-rabbit IgG (Invitrogen, Carlsbad, CA; diluted to 1:100) and Coumarin AMCA donkey anti-human IgG (Jackson ImmunoResearch, West Grove, PA; diluted to 1:200) were suspended in ADB. The last incubation of Alexa Fluor 568 donkey anti-goat (Invitrogen, Carlsbad, CA; diluted 1:100) was incubated at 1:100 for 2 hours at 37* C. Slides were fixed with Prolong Gold Antifade (Invitrogen, Carlsbad, CA) for 24 hours after a final wash in 1x PBS. Three slides of cell spreads per mouse were prepared to serve as technical replicates for the staining protocol. Comparisons of multiple, stained slides from the same mouse showed no difference in mean MLH1 cell counts and mean cell quality. Sampled numbers of mice and cells per mouse were maximized to the extent possible given constraints on breeding and time.

### Image Processing

Images were captured using a Zeiss Axioplan 2 microscope with AxioLab camera and AxioVision software (Zeiss, Cambridge, UK). The number of cells imaged per individual mouse is based on previous studies (Dumont and Payseur, 2011; Murdoch et al., 2010; Wang and Payseur, 2017). Preprocessing, including cropping, noise reduction, and histogram adjustments, was performed using Photoshop (v13.0). Image file names were anonymized before manual scoring of MLH1 foci or DMC1 foci using Photoshop.

### Analyses

To estimate the number of crossovers across the genome, we counted MLH1 foci within bivalents, synapsed homologous chromosomes. MLH1 foci were counted in pachytene cells with intact and complete karyotypes (19 acrocentric bivalents and XY for spermatocytes; 20 acrocentric bivalents for oocytes) and distinct MLH1 foci. A quality score ranging from 1 (best) to 5 (worst) was assigned to each cell based on visual appearance of staining and spread of bivalents. Cells with a score of 5 were excluded from the final analysis. Distributions of MLH1 count per cell were visually inspected for normality (Supplemental Figure 1). When outliers for MLH1 count were found during preliminary analysis, the original images were inspected and the counts confirmed.

MLH1 foci located on the XY in spermatocytes were excluded from counts. In addition to MLH1 counts, we measured several traits to further characterize the recombination landscape. To estimate the number of double-strand breaks, a minority of which lead to crossovers, mean DMC1 foci per cell was quantified for a single male from each of a subset of strains (*molossinus^MSM^*, *musculus^PWD^*, *domesticus^WSB^*, and *domesticus^G^*). SC morphology and CREST foci number were used to stage spermatocytes as early zygotene or late zygotene.

To measure bivalent SC length, two image analysis algorithms were used. The first algorithm estimates the total (summed) SC length across bivalents for individual cells (Wang et al., 2019). The second algorithm estimates the SC length of individual bivalents (Peterson et al., 2019). Both algorithms apply a ‘skeletonizing’ transformation to synapsed chromosomes that produces a single, pixel-wide ‘trace’ of the bivalent shape. Total SC length per cell was quantified from pachytene cell images (Wang et al., 2019).

To reduce algorithmic errors in SC isolation, outliers were visually identified at the mouse level and removed from the data set. Mouse averages were calculated from cell-wide total SC lengths in 3,195 out of 3,871 cells with MLH1 counts. SC length of individual bivalents was quantified in pachytene cell images (Peterson et al., 2019). The DNA CrossOver algorithm (Peterson et al., 2019) isolates single, straightened bivalent shapes, returning SC length, location of MLH1 foci, and location of CREST (centromere) foci. The algorithm substantially speeds the accurate measurement of bivalents, but it sometimes interprets overlapping bivalents as single bivalents. In our data set, average proportions of bivalents per cell isolated by the algorithm ranged from 0.48 (*molossinus^MSM^* male) to 0.72 (*musculus^KAZ^* female). From the total set of pachytene cell images, 10,213 bivalent objects were isolated by the algorithm. Following a manual curation, 9,569 single-bivalent observations remained. The accuracy of the algorithm is high compared to hand measures after this curation step (Peterson et al., 2019). The curated single bivalent data supplemented our cell-wide MLH1 count data with MLH1 foci counts for single bivalents. Proportions of bivalents with the same number of MLH1 foci were compared across strains using a chi-square test.

To account for confounding effects of sex chromosomes from pooled samples of bivalents, we also considered a reduced data set including only bivalents with SC lengths below the 2nd quartile in cells with at least 17 of 20 single bivalent measures. This “short bivalent” data set included the four or five shortest bivalents within a cell, thus excluding the X bivalent in oocytes. A total of 699 short bivalents were isolated from 102 oocytes and 42 spermatocytes. Although this smaller data set had decreased power, it offered a more comparable set of single bivalents to compare between the sexes. A “long bivalent” data set was formed from those bivalents above the 4th quartile in SC lengths per cell. A total of 703 long bivalents were isolated from 102 oocytes and 42 spermatocytes.

To examine crossover interference, the distance (in SC units) between MLH1 foci (inter-focal distance; IFD_raw_) was measured for those single bivalents containing two MLH1 foci. A normalized measure of interference (IFD_norm_) was computed by dividing IFD_raw_ by SC length on a per-bivalent basis.

We used a series of statistical models to interpret patterns of variation in the recombination traits we measured (Table 2). We used mouse average as the dependent variable in all analyses. We first constructed a linear mixed model (M1) using lmer() from the lmer4 package (Bates et al., 2015) in R (v3.5.2) (Team, 2015). In this model, strain was coded as a random effect, with significance evaluated using a likelihood ratio test using exactRLRT() from RLRsim (Scheipl et al., 2008). Subspecies, sex, and their interaction were coded as fixed effects, with significance evaluated using a chi-square test comparing the full and reduced models (drop1() and anova()) (Bates et al., 2015). The hierarchical nature of the data meant that nesting of levels across observations was implicit (*i.e.* mouse within strain, within subspecies) and not explicitly coded. We used the subspecies effect to quantify divergence between subspecies and the (random) strain effect to quantify variation within subspecies in a sex-specific manner. In separate analyses using model M1, we considered mouse averages as dependent variables for each of the following traits: MLH1 count per cell, total SC length per cell, single bivalent SC length per cell, IFD_raw_, IFD_norm_, and average MLH1 position (for single-focus bivalents). Four additional linear models containing only fixed effects (M2-M5) (Table 2) were used to further investigate results obtained from model M1.

## Acknowledgements

This research was funded by NIH grants R01GM120051 and R01GM100426 to B. A. P.. A. L. P. was partly supported by NIH T32GM007133. We are grateful to Francisco Pelegri for generous assistance with microscopy. We thank Karl Broman and Cécile Ané for advice on analyses.

## Supplemental Figures

**Supplemental Figure 1.**
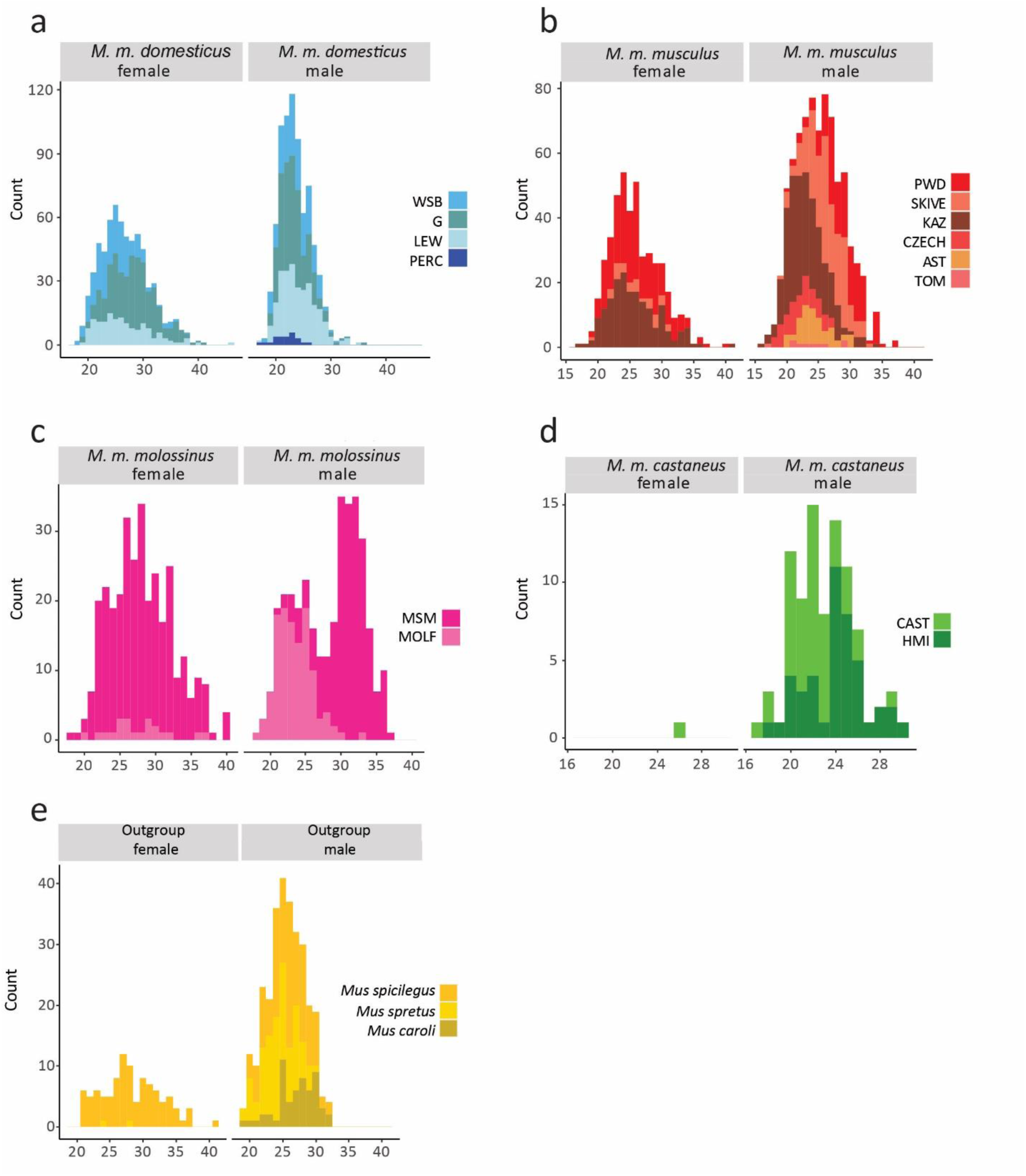
Distributions of MLH1 Counts per Cell. Strain names are abbreviated for space.

**Supplemental Figure 2.**
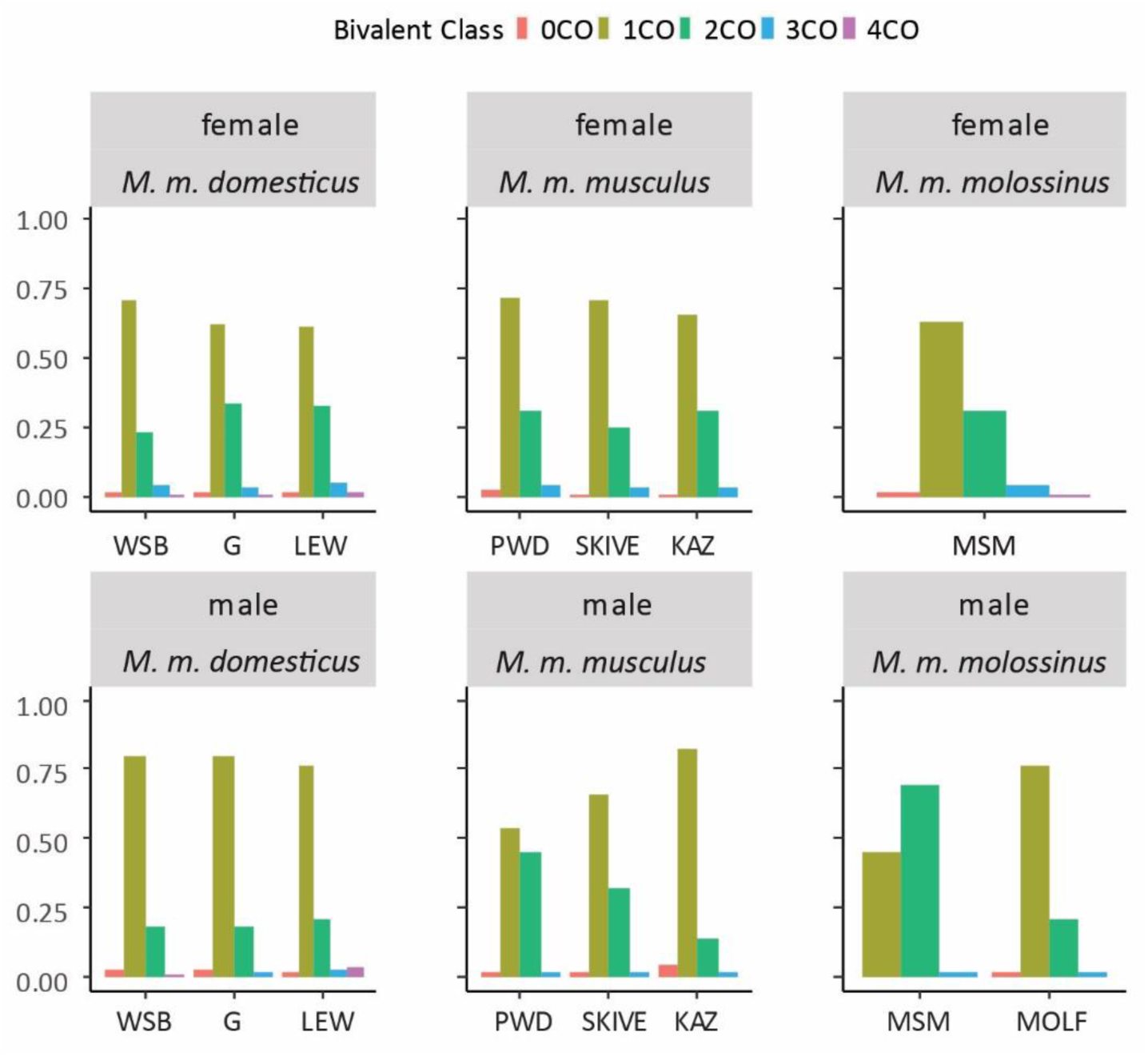
Proportions of Bivalents with Different Numbers of MLH1 Foci. Strain names are abbreviated for space.

**Supplemental Figure 3.**
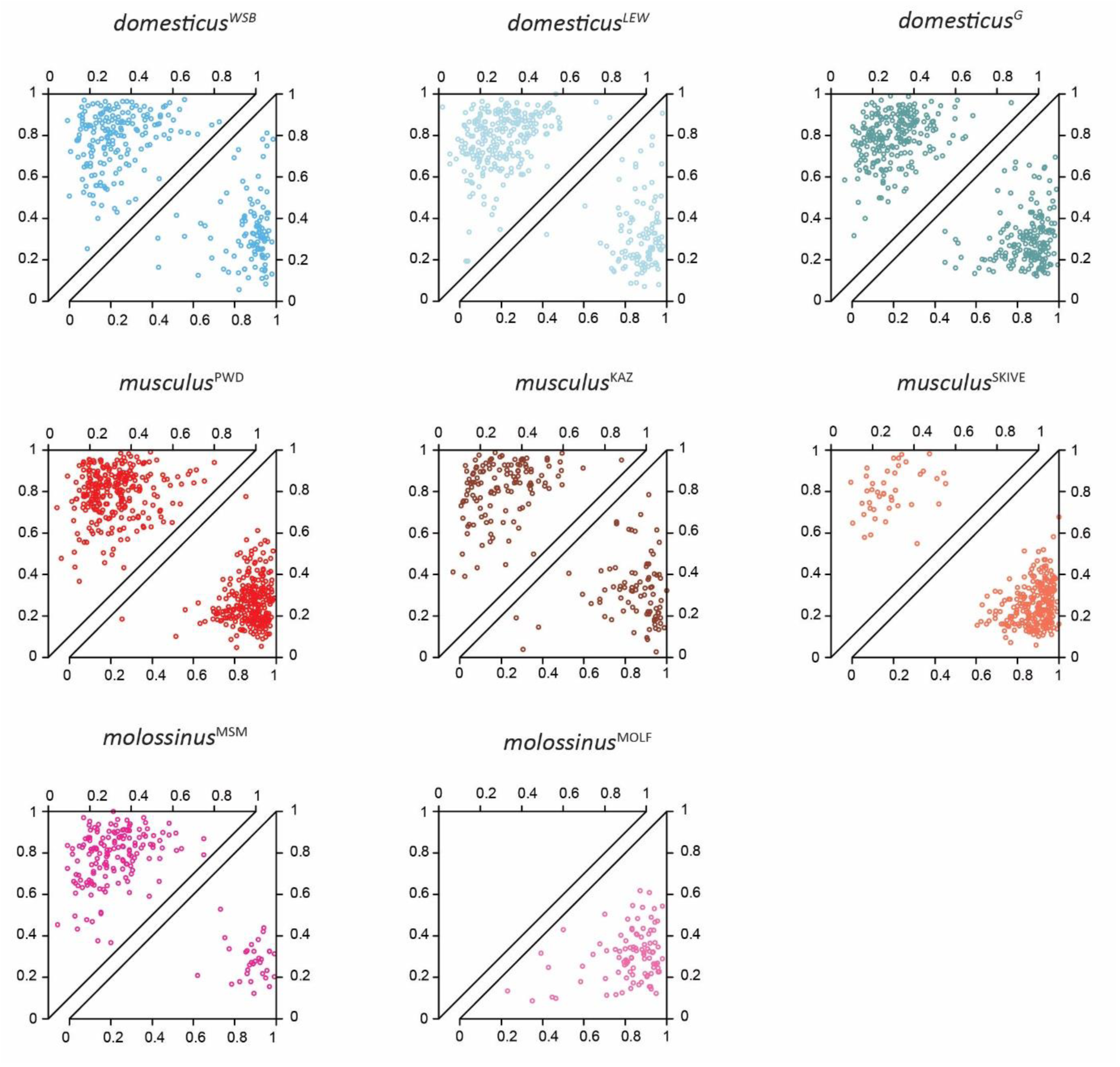
Inter-focal Distances on Bivalents with Two MLH1 Foci. Each point shows the positions of both foci, normalized by bivalent SC length. Observations are separated by sex (females=top triangles; males=bottom triangles).

## Supplemental Tables

**Supplemental Table 1.**

| **M1 MLH1 Count** | **p values** |  | **Coefficients**<br>**(fixed estimates)** | **Random Effects**<br>**(standard deviation)** |
| --- | --- | --- | --- | --- |
| Subspecies | 0.00097 | Intercept | 26.356 |  |
| Sex | 0.00000 | Subspecies Musculus | -0.755 |  |
| Subspecies*Sex | 0.00018 | Subspecies Molossinus | -0.482 |  |
| strain(random) | 0.00010 | Sex(male) | -2.649 |  |
|  |  | Musculus*male | 2.953 |  |
|  |  | Molossinus*male | 3.201 |  |
|  |  | intercept |  | 1.69 |
|  |  | Strain |  | 1.89 |

**Supplemental Table 2.**

| **M2 MLH1 Count** | **Estimate** | **Std. Error** | **t value** | **Pr(>\|t\|)** |
| --- | --- | --- | --- | --- |
| (Intercept) | 24.718 | 0.447 | 55.356 | 0.000 |
| Subspecies Musculus | 0.849 | 0.714 | 1.190 | 0.236 |
| Subspecies Molossinus | 2.901 | 1.729 | 1.678 | 0.096 |
| Sex (male) | -1.194 | 0.692 | -1.726 | 0.087 |
| Strain G | 3.301 | 0.657 | 5.023 | 0.000 |
| Strain LEW | 1.694 | 0.714 | 2.373 | 0.019 |
| Strain PWD | 0.257 | 0.704 | 0.365 | 0.716 |
| Strain MSM | 0.086 | 1.729 | 0.050 | 0.960 |
| Strain SKIVE | 0.371 | 1.761 | 0.210 | 0.834 |
| Subspecies Musculus * Sex | -0.768 | 1.021 | -0.753 | 0.453 |
| Subspecies Molossinus * Sex | -3.185 | 1.933 | -1.648 | 0.102 |
| Strain G * Sex (male) | -3.144 | 0.982 | -3.201 | 0.002 |
| Strain LEW * Sex (male) | -1.165 | 1.090 | -1.070 | 0.287 |
| Strain PWD * Sex (male) | 4.444 | 1.048 | 4.239 | 0.000 |
| Strain MSM * Sex (male) | 6.826 | 2.038 | 3.349 | 0.001 |
| Strain SKIVE * Sex (male) | 2.260 | 1.978 | 1.143 | 0.255 |

**Supplemental Table 3.**
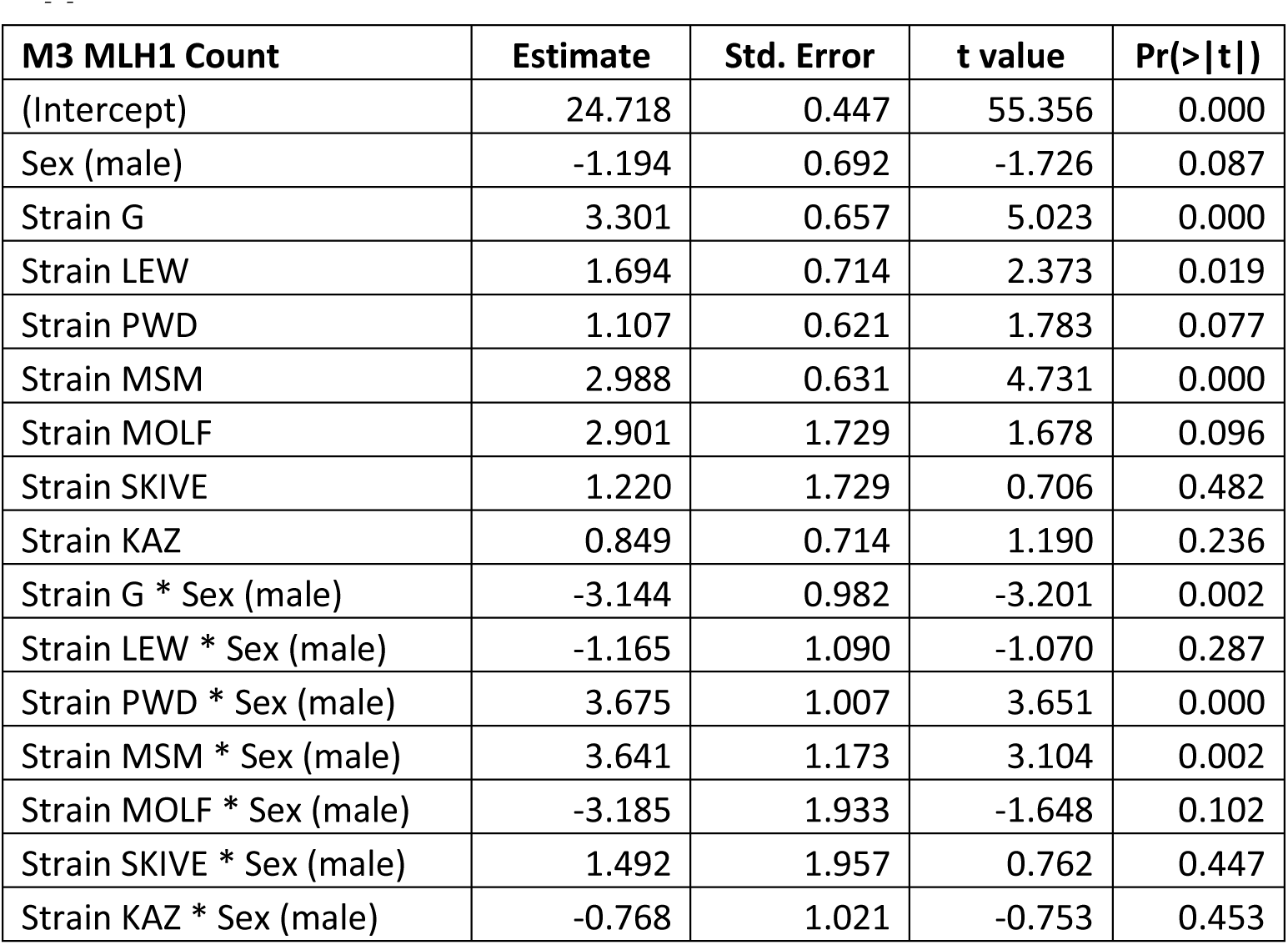

**Supplemental Table 4.**
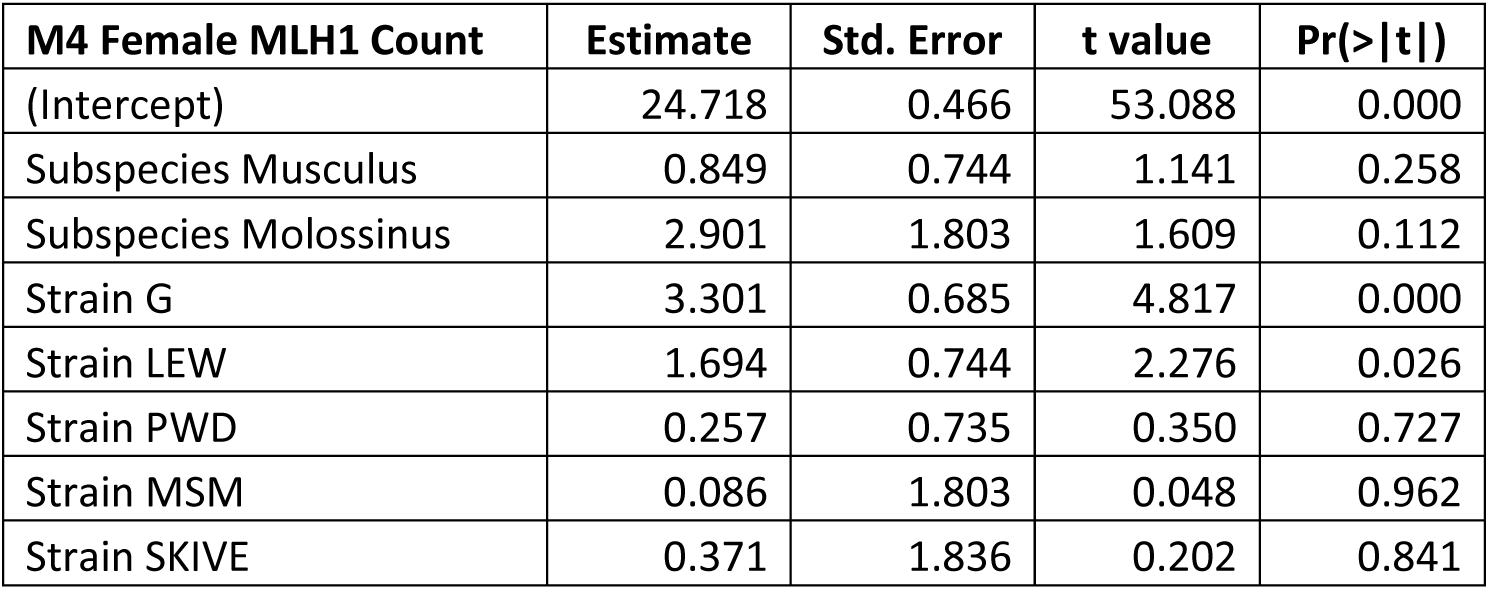

| **M4 Female MLH1 Count** | **Estimate** | **Std. Error** | **t value** | **Pr(>\|t\|)** |
| --- | --- | --- | --- | --- |
| (Intercept) | 24.718 | 0.466 | 53.088 | 0.000 |
| Subspecies Musculus | 0.849 | 0.744 | 1.141 | 0.258 |
| Subspecies Molossinus | 2.901 | 1.803 | 1.609 | 0.112 |
| Strain G | 3.301 | 0.685 | 4.817 | 0.000 |
| Strain LEW | 1.694 | 0.744 | 2.276 | 0.026 |
| Strain PWD | 0.257 | 0.735 | 0.350 | 0.727 |
| Strain MSM | 0.086 | 1.803 | 0.048 | 0.962 |
| Strain SKIVE | 0.371 | 1.836 | 0.202 | 0.841 |

**Supplemental Table 5.**
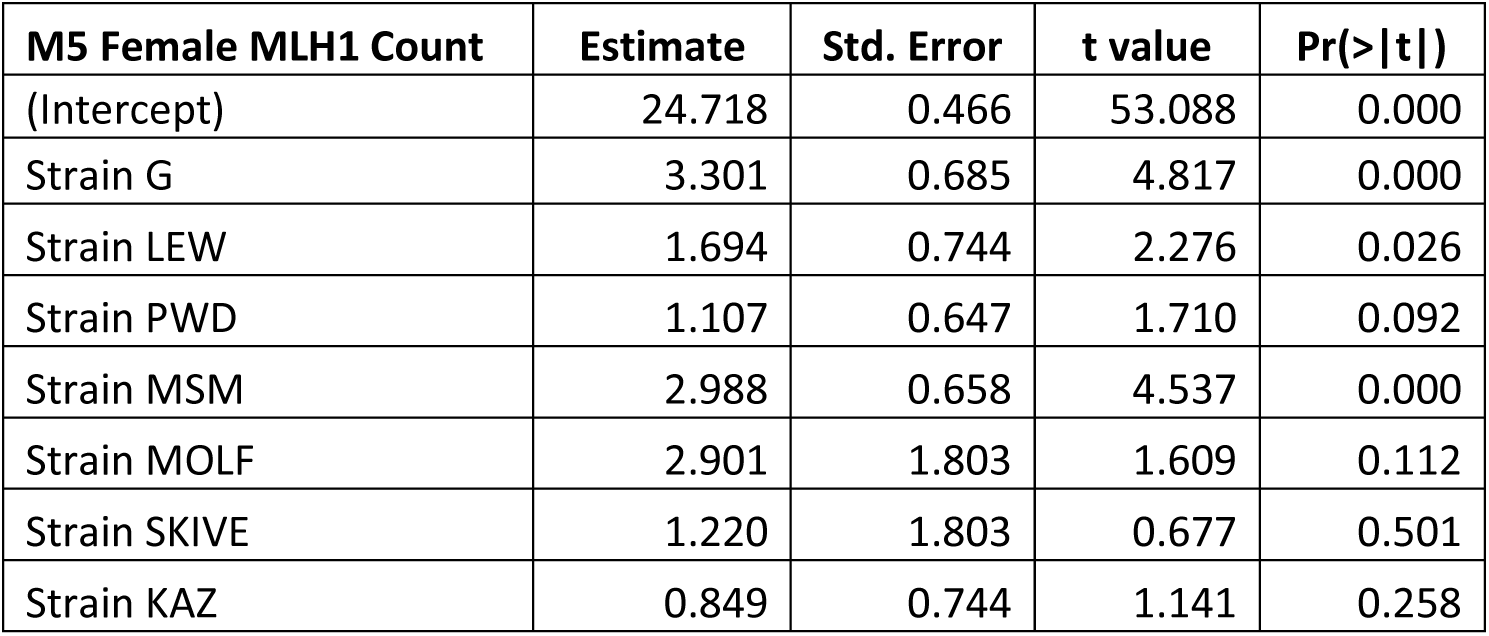

**Supplemental Table 6.**

| **M4 Male MLH1 Count** | **Estimate** | **Std. Error** | **t value** | **Pr(>\|t\|)** |
| --- | --- | --- | --- | --- |
| (Intercept) | 23.524 | 0.495 | 47.530 | 0.000 |
| Subspecies Musculus | -1.330 | 1.030 | -1.291 | 0.202 |
| Subspecies Molossinus | -0.284 | 0.808 | -0.352 | 0.727 |
| Strain G | 0.157 | 0.684 | 0.229 | 0.820 |
| Strain LEW | 0.528 | 0.771 | 0.685 | 0.496 |
| Strain PERC | -1.716 | 1.641 | -1.045 | 0.300 |
| Strain PWD | 6.113 | 1.060 | 5.769 | 0.000 |
| Strain MSM | 6.913 | 1.010 | 6.843 | 0.000 |
| Strain SKIVE | 4.042 | 1.143 | 3.537 | 0.001 |
| Strain KAZ | 1.411 | 1.019 | 1.385 | 0.172 |
| Strain TOM | 3.406 | 1.807 | 1.885 | 0.065 |
| Strain AST | 2.703 | 1.807 | 1.496 | 0.140 |

**Supplemental Table 7.**

| **M5 Male MLH1 Count** | **Estimate** | **Std. Error** | **t value** | **Pr(>\|t\|)** |
| --- | --- | --- | --- | --- |
| (Intercept) | 23.524 | 0.495 | 47.530 | 0.000 |
| Strain G | 0.157 | 0.684 | 0.229 | 0.820 |
| Strain LEW | 0.528 | 0.771 | 0.685 | 0.496 |
| Strain PERC | -1.716 | 1.641 | -1.045 | 0.300 |
| Strain PWD | 4.782 | 0.742 | 6.442 | 0.000 |
| Strain MSM | 6.629 | 0.926 | 7.159 | 0.000 |
| Strain MOLF | -0.284 | 0.808 | -0.352 | 0.727 |
| Strain SKIVE | 2.712 | 0.857 | 3.164 | 0.003 |
| Strain KAZ | 0.081 | 0.684 | 0.118 | 0.906 |
| Strain TOM | 2.076 | 1.641 | 1.265 | 0.211 |
| Strain AST | 1.373 | 1.641 | 0.836 | 0.406 |
| Strain CZECH | -1.330 | 1.030 | -1.291 | 0.202 |

**Supplemental Table 8.**

| **M1 Total SC Length** | **p values** |  | **Coefficients**<br>**(fixed estimates)** | **Random Effects**<br>**(standard deviation)** |
| --- | --- | --- | --- | --- |
| Subspecies | 0.00085 | Intercept | 1960.2 |  |
| Sex | 0.00000 | Subspecies Musculus | -44.1 |  |
| Subspecies*Sex | 0.00010 | Subspecies Molossinus | -119.1 |  |
| Strain(random) | 0.00010 | Sex(male) | -558 |  |
|  |  | Musculus*male | 167.1 |  |
|  |  | Molossinus*male | 396.9 |  |
|  |  | Intercept |  | 119 |
|  |  | Strain |  | 263 |

**Supplemental Table 9.**

| **M2 Total SC Length** | **Estimate** | **Std. Error** | **t value** | **Pr(>\|t\|)** |
| --- | --- | --- | --- | --- |
| (Intercept) | 1683.383 | 36.479 | 46.147 | 0.000 |
| Subspecies Musculus | 229.568 | 62.002 | 3.703 | 0.000 |
| Subspecies Molossinus | 161.137 | 53.281 | 3.024 | 0.003 |
| Strain G | 431.238 | 54.282 | 7.944 | 0.000 |
| Strain LEW | 248.684 | 59.941 | 4.149 | 0.000 |
| Strain PWD | -26.764 | 61.403 | -0.436 | 0.664 |
| Strain SKIVE | 50.076 | 90.383 | 0.554 | 0.580 |
| Sex (male) | -345.005 | 58.200 | -5.928 | 0.000 |
| Subspecies Musculus * Sex (male) | -84.338 | 89.204 | -0.945 | 0.346 |
| Subspecies Molossinus * Sex (male) | 243.125 | 97.055 | 2.505 | 0.013 |
| Strain G * Sex (male) | -303.270 | 84.021 | -3.609 | 0.000 |
| Strain LEW * Sex (male) | -169.848 | 94.240 | -1.802 | 0.074 |
| Strain PWD * Sex (male) | 121.505 | 93.030 | 1.306 | 0.194 |
| Strain SKIVE * Sex (male) | -119.766 | 121.449 | -0.986 | 0.326 |

**Supplemental Table 10.**

| **M3 Total SC Length** | **Estimate** | **Std. Error** | **t value** | **Pr(>\|t\|)** |
| --- | --- | --- | --- | --- |
| (Intercept) | 1683.383 | 36.479 | 46.147 | 0.000 |
| Strain G | 431.238 | 54.282 | 7.944 | 0.000 |
| Strain LEW | 248.684 | 59.941 | 4.149 | 0.000 |
| Strain PWD | 202.805 | 50.867 | 3.987 | 0.000 |
| Strain MSM | 161.137 | 53.281 | 3.024 | 0.003 |
| Strain SKIVE | 279.644 | 83.583 | 3.346 | 0.001 |
| Strain KAZ | 229.568 | 62.002 | 3.703 | 0.000 |
| Sex (male) | -345.005 | 58.200 | -5.928 | 0.000 |
| Strain G * Sex (male) | -303.270 | 84.021 | -3.609 | 0.000 |
| Strain LEW * Sex (male) | -169.848 | 94.240 | -1.802 | 0.074 |
| Strain PWD * Sex (male) | 37.168 | 86.439 | 0.430 | 0.668 |
| Strain MSM * Sex (male) | 243.125 | 97.055 | 2.505 | 0.013 |
| Strain SKIVE * Sex (male) | -204.104 | 116.478 | -1.752 | 0.082 |
| Strain KAZ * Sex (male) | -84.338 | 89.204 | -0.945 | 0.346 |

**Supplemental Table 11.**
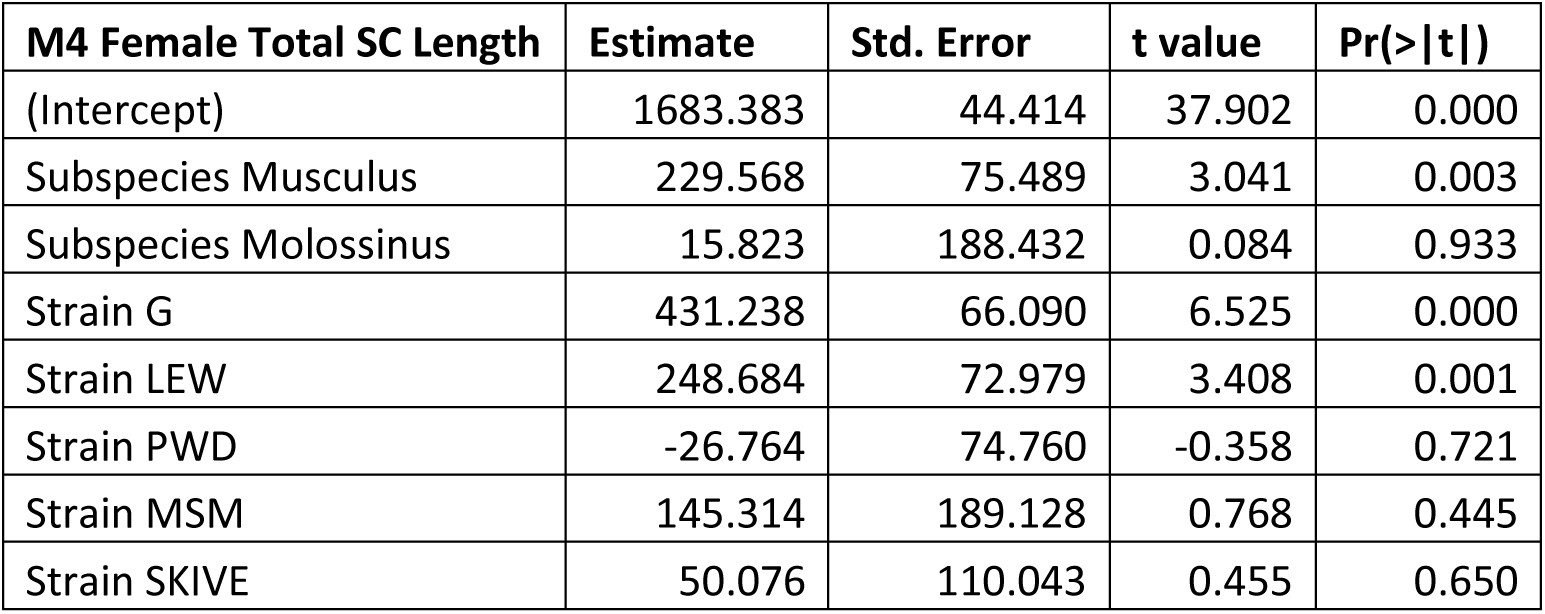

**Supplemental Table 12.**

| **M5 Female Total SC Length** | **Estimate** | **Std. Error** | **t value** | **Pr(>\|t\|)** |
| --- | --- | --- | --- | --- |
| (Intercept) | 1683.383 | 44.414 | 37.902 | 0.000 |
| Strain G | 431.238 | 66.090 | 6.525 | 0.000 |
| Strain LEW | 248.684 | 72.979 | 3.408 | 0.001 |
| Strain PWD | 202.805 | 61.932 | 3.275 | 0.002 |
| Strain MSM | 161.137 | 64.870 | 2.484 | 0.015 |
| Strain MOLF | 15.823 | 188.432 | 0.084 | 0.933 |
| Strain SKIVE | 279.644 | 101.765 | 2.748 | 0.007 |
| Strain KAZ | 229.568 | 75.489 | 3.041 | 0.003 |

**Supplemental Table 13.**

| **M4 Male Total SC Length** | **Estimate** | **Std. Error** | **t value** | **Pr(>\|t\|)** |
| --- | --- | --- | --- | --- |
| (Intercept) | 1338.379 | 23.387 | 57.228 | 0.000 |
| Subspecies Musculus | 236.879 | 41.835 | 5.662 | 0.000 |
| Subspecies Molossinus | 213.996 | 37.502 | 5.706 | 0.000 |
| Strain G | 127.967 | 33.074 | 3.869 | 0.000 |
| Strain LEW | 78.836 | 37.502 | 2.102 | 0.040 |
| Strain PERC | 109.979 | 81.014 | 1.358 | 0.179 |
| Strain PWD | 3.093 | 44.219 | 0.070 | 0.944 |
| Strain MSM | 190.266 | 45.417 | 4.189 | 0.000 |
| Strain SKIVE | -161.339 | 49.056 | -3.289 | 0.002 |
| Strain KAZ | -91.648 | 41.835 | -2.191 | 0.032 |
| Strain TOM | 72.493 | 64.895 | 1.117 | 0.268 |
| Strain AST | 39.557 | 64.895 | 0.610 | 0.544 |

**Supplemental Table 14.**

| **M5 Male Total SC Length** | **Estimate** | **Std. Error** | **t value** | **Pr(>\|t\|)** |
| --- | --- | --- | --- | --- |
| (Intercept) | 1338.379 | 23.387 | 57.228 | 0.000 |
| Strain G | 127.967 | 33.074 | 3.869 | 0.000 |
| Strain LEW | 78.836 | 37.502 | 2.102 | 0.040 |
| Strain PERC | 109.979 | 81.014 | 1.358 | 0.179 |
| Strain PWD | 239.972 | 36.041 | 6.658 | 0.000 |
| Strain MSM | 404.262 | 41.835 | 9.663 | 0.000 |
| Strain MOLF | 213.996 | 37.502 | 5.706 | 0.000 |
| Strain SKIVE | 75.540 | 41.835 | 1.806 | 0.076 |
| Strain KAZ | 145.230 | 33.074 | 4.391 | 0.000 |
| Strain TOM | 309.371 | 59.625 | 5.189 | 0.000 |
| Strain AST | 276.436 | 59.625 | 4.636 | 0.000 |
| Strain CZECH | 236.879 | 41.835 | 5.662 | 0.000 |

**Supplemental Table 15.**

| **M1 Short Bivalent SC Length** | **p values** |  | **Coefficients (fixed estimates)** | **Random Effects (standard deviation)** |
| --- | --- | --- | --- | --- |
| Subspecies | 0.12684 | Intercept | 80.34 |  |
| Sex | 0.00000 | Subspecies Musculus | -5.71 |  |
| Subspecies*Sex | 0.02989 | Subspecies Molossinus | -4.65 |  |
| Strain(random) | 0.18840 | Sex(male) | -26.26 |  |
|  |  | Musculus*male | 10.52 |  |
|  |  | Molossinus*male |  |  |
|  |  | Intercept |  | 2.49 |
|  |  | Strain |  | 8.01 |

**Supplemental Table 16.**

| **M2 Short Bivalent SC Length** | **Estimate** | **Std. Error** | **t value** | **Pr(>\|t\|)** |
| --- | --- | --- | --- | --- |
| (Intercept) | 73.886 | 4.633 | 15.947 | 0.000 |
| Subspecies Musculus | 6.897 | 6.129 | 1.125 | 0.267 |
| Subspecies Molossinus | 1.803 | 6.552 | 0.275 | 0.785 |
| Strain G | 7.965 | 5.674 | 1.404 | 0.168 |
| Strain LEW | 9.166 | 5.674 | 1.615 | 0.114 |
| Strain PWD | -5.068 | 4.914 | -1.031 | 0.309 |
| Strain SKIVE | -13.897 | 5.674 | -2.449 | 0.019 |
| Sex (male) | -25.336 | 7.326 | -3.459 | 0.001 |
| Subspecies Musculus * Sex (male) | 1.960 | 9.551 | 0.205 | 0.838 |
| Strain G * Sex (male) | 0.100 | 8.972 | 0.011 | 0.991 |
| Strain LEW * Sex (male) | -2.216 | 9.828 | -0.226 | 0.823 |
| Strain PWD * Sex (male) | 7.727 | 8.190 | 0.943 | 0.351 |
| Strain SKIVE * Sex (male) | 15.180 | 8.352 | 1.817 | 0.077 |

**Supplemental Table 17.**

| **M3 Short Bivalent SC Length** | **Estimate** | **Std. Error** | **t value** | **Pr(>\|t\|)** |
| --- | --- | --- | --- | --- |
| (Intercept) | 73.886 | 4.633 | 15.947 | 0.000 |
| Strain G | 7.965 | 5.674 | 1.404 | 0.168 |
| Strain LEW | 9.167 | 5.674 | 1.615 | 0.114 |
| Strain PWD | 1.829 | 5.433 | 0.337 | 0.738 |
| Strain MSM | 1.803 | 6.552 | 0.275 | 0.785 |
| Strain SKIVE | -7.000 | 6.129 | -1.142 | 0.260 |
| Strain KAZ | 6.897 | 6.129 | 1.125 | 0.267 |
| Sex (male) | -25.336 | 7.326 | -3.459 | 0.001 |
| Strain G * Sex (male) | 0.100 | 8.972 | 0.011 | 0.991 |
| Strain LEW * Sex (male) | -2.217 | 9.828 | -0.226 | 0.823 |
| Strain PWD * Sex (male) | 9.687 | 9.120 | 1.062 | 0.295 |
| Strain SKIVE * Sex (male) | 17.140 | 9.266 | 1.850 | 0.072 |
| Strain KAZ * Sex (male) | 1.960 | 9.551 | 0.205 | 0.838 |

**Supplemental Table 18.**

| **M4 Female Short Bivalent**<br>**SC Length** | **Estimate** | **Std. Error** | **t value** | **Pr(>\|t\|)** |
| --- | --- | --- | --- | --- |
| (Intercept) | 73.886 | 5.334 | 13.853 | 0.000 |
| Subspecies Musculus | 6.897 | 7.056 | 0.977 | 0.337 |
| Subspecies Molossinus | 1.803 | 7.543 | 0.239 | 0.813 |
| Strain G | 7.965 | 6.532 | 1.219 | 0.233 |
| Strain LEW | 9.167 | 6.532 | 1.403 | 0.172 |
| Strain PWD | -5.068 | 5.657 | -0.896 | 0.378 |
| Strain SKIVE | -13.897 | 6.532 | -2.127 | 0.043 |

**Supplemental Table 19.**

| **M5 Female Short Bivalent**<br>**SC Length** | **Estimate** | **Std. Error** | **t value** | **Pr(>\|t\|)** |
| --- | --- | --- | --- | --- |
| (Intercept) | 73.886 | 5.334 | 13.853 | 0.000 |
| Strain G | 7.965 | 6.532 | 1.219 | 0.233 |
| Strain LEW | 9.167 | 6.532 | 1.403 | 0.172 |
| Strain PWD | 1.829 | 6.254 | 0.292 | 0.772 |
| Strain MSM | 1.803 | 7.543 | 0.239 | 0.813 |
| Strain SKIVE | -7.000 | 7.056 | -0.992 | 0.330 |
| Strain KAZ | 6.897 | 7.056 | 0.977 | 0.337 |

**Supplemental Table 20.**

| **M4 Male Short Bivalent**<br>**SC Length** | **Estimate** | **Std. Error** | **t value** | **Pr(>\|t\|)** |
| --- | --- | --- | --- | --- |
| (Intercept) | 48.550 | 3.120 | 15.563 | 0.000 |
| Subspecies Musculus | 12.992 | 4.412 | 2.945 | 0.011 |
| Subspecies Molossinus | 13.762 | 5.403 | 2.547 | 0.024 |
| Strain G | 8.065 | 3.821 | 2.111 | 0.055 |
| Strain LEW | 6.950 | 4.412 | 1.575 | 0.139 |
| Strain PWD | -1.475 | 4.027 | -0.366 | 0.720 |
| Strain SKIVE | -2.852 | 3.821 | -0.746 | 0.469 |
| Strain KAZ | -4.135 | 4.027 | -1.027 | 0.323 |

**Supplemental Table 21.**
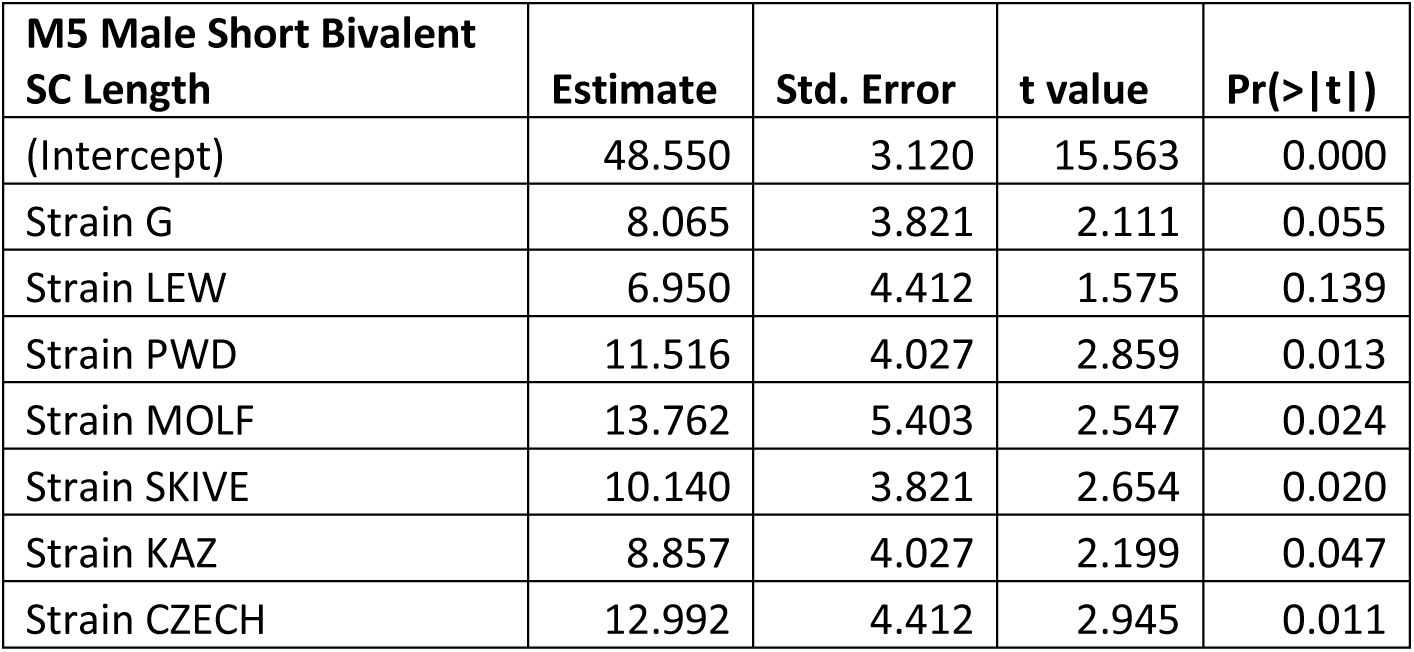

**Supplemental Table 22.**

| **M4 Male Long Bivalent**<br>**SC Length** | **Estimate** | **Std. Error** | **t value** | **Pr(>\|t\|)** |
| --- | --- | --- | --- | --- |
| (Intercept) | 86.400 | 4.942 | 17.482 | 0.000 |
| Subspecies Musculus | 17.836 | 6.989 | 2.552 | 0.024 |
| Subspecies Molossinus | 11.457 | 8.560 | 1.338 | 0.204 |
| Strain G | 9.469 | 6.053 | 1.564 | 0.142 |
| Strain LEW | 9.000 | 6.989 | 1.288 | 0.220 |
| Strain PWD | -3.290 | 6.380 | -0.516 | 0.615 |
| Strain SKIVE | -4.205 | 6.053 | -0.695 | 0.499 |
| Strain KAZ | -5.837 | 6.380 | -0.915 | 0.377 |

**Supplemental Table 23.**

| **M5 Male Long Bivalent**<br>**SC Length** | **Estimate** | **Std. Error** | **t value** | **Pr(>\|t\|)** |
| --- | --- | --- | --- | --- |
| (Intercept) | 86.400 | 4.942 | 17.482 | 0.000 |
| Strain G | 9.469 | 6.053 | 1.564 | 0.142 |
| Strain LEW | 9.000 | 6.989 | 1.288 | 0.220 |
| Strain PWD | 14.546 | 6.380 | 2.280 | 0.040 |
| Strain MOLF | 11.457 | 8.560 | 1.338 | 0.204 |
| Strain SKIVE | 13.631 | 6.053 | 2.252 | 0.042 |
| Strain KAZ | 11.999 | 6.380 | 1.881 | 0.083 |
| Strain CZECH | 17.836 | 6.989 | 2.552 | 0.024 |

**Supplemental Table 24.**

| **M1 Normalized Foci Position** | **p values** |  | **Coefficients (fixed estimates)** | **Random Effects (standard deviation)** |
| --- | --- | --- | --- | --- |
| Subspecies | 0.124 | Intercept | 0.559 |  |
| Sex | 0.000 | Subspecies Musculus | 0.009 |  |
| Subspecies*Sex | 0.056 | Subspecies Molossinus | 0.016 |  |
| Strain(random) | 0.003 | Sex(male) | 0.137 |  |
|  |  | Musculus*male | -0.031 |  |
|  |  | Molosinus*male | 0.020 |  |
|  |  | Intercept |  | 0.019 |
|  |  | Strain |  | 0.031 |

**Supplemental Table 25.**

| **M2 Normalized Foci Position** | **Estimate** | **Std. Error** | **t value** | **Pr(>\|t\|)** |
| --- | --- | --- | --- | --- |
| (Intercept) | 0.590 | 0.018 | 32.808 | 0.000 |
| Subspecies Musculus | -0.043 | 0.023 | -1.907 | 0.061 |
| Subspecies Molossinus | -0.016 | 0.024 | -0.659 | 0.512 |
| Strain G | -0.039 | 0.022 | -1.753 | 0.085 |
| Strain LEW | -0.051 | 0.021 | -2.406 | 0.019 |
| Strain PWD | 0.028 | 0.017 | 1.653 | 0.103 |
| Strain SKIVE | 0.030 | 0.021 | 1.440 | 0.155 |
| Sex (male) | 0.142 | 0.023 | 6.251 | 0.000 |
| Subspecies Musculus * Sex (male) | -0.010 | 0.032 | -0.313 | 0.755 |
| Subspecies Molossinus * Sex (male) | 0.014 | 0.035 | 0.402 | 0.689 |
| Strain G * Sex (male) | -0.025 | 0.028 | -0.876 | 0.385 |
| Strain LEW * Sex (male) | 0.010 | 0.029 | 0.360 | 0.720 |
| Strain PWD * Sex (male) | -0.040 | 0.028 | -1.409 | 0.164 |
| Strain SKIVE * Sex (male) | -0.030 | 0.030 | -1.007 | 0.318 |

**Supplemental Table 26.**

| **M3 Normalized Foci Position** | **Estimate** | **Std. Error** | **t value** | **Pr(>\|t\|)** |
| --- | --- | --- | --- | --- |
| (Intercept) | 0.590 | 0.018 | 32.808 | 0.000 |
| Strain G | -0.039 | 0.022 | -1.753 | 0.085 |
| Strain LEW | -0.051 | 0.021 | -2.406 | 0.019 |
| Strain PWD | -0.016 | 0.020 | -0.769 | 0.445 |
| Strain MSM | -0.016 | 0.024 | -0.659 | 0.512 |
| Strain SKIVE | -0.013 | 0.024 | -0.559 | 0.578 |
| Strain KAZ | -0.043 | 0.023 | -1.907 | 0.061 |
| Sex (male) | 0.142 | 0.023 | 6.251 | 0.000 |
| Strain G * Sex (male) | -0.025 | 0.028 | -0.876 | 0.385 |
| Strain LEW * Sex (male) | 0.010 | 0.029 | 0.360 | 0.720 |
| Strain PWD * Sex (male) | -0.050 | 0.028 | -1.765 | 0.082 |
| Strain MSM * Sex (male) | 0.014 | 0.035 | 0.402 | 0.689 |
| Strain SKIVE * Sex (male) | -0.040 | 0.030 | -1.343 | 0.184 |
| Strain KAZ * Sex (male) | -0.010 | 0.032 | -0.313 | 0.755 |

**Supplemental Table 27.**

| **M4 Female**<br>**Normalized Foci Position** | **Estimate** | **Std. Error** | **t value** | **Pr(>\|t\|)** |
| --- | --- | --- | --- | --- |
| (Intercept) | 0.507 | 0.008 | 64.388 | 0.000 |
| Subspecies Musculus | -0.007 | 0.013 | -0.573 | 0.566 |
| Subspecies Molossinus | -0.017 | 0.012 | -1.407 | 0.160 |
| Strain G | -0.044 | 0.011 | -3.891 | 0.000 |
| Strain LEW | -0.063 | 0.011 | -5.466 | 0.000 |
| Strain PWD | -0.010 | 0.012 | -0.868 | 0.385 |
| Strain SKIVE | -0.003 | 0.014 | -0.231 | 0.817 |

**Supplemental Table 28.**
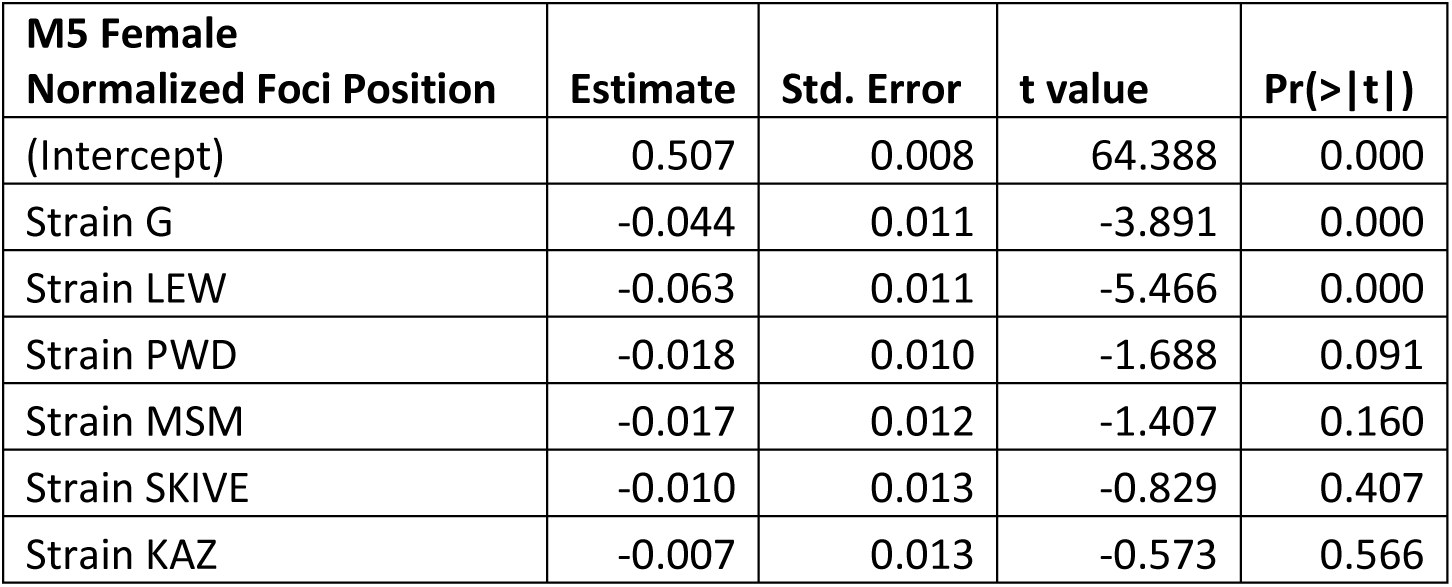

**Supplemental Table 29.**

| **M4 Male**<br>**Normalized Foci Position** | **Estimate** | **Std. Error** | **t value** | **Pr(>\|t\|)** |
| --- | --- | --- | --- | --- |
| (Intercept) | 0.733 | 0.014 | 52.208 | 0.000 |
| Subspecies Musculus | -0.042 | 0.023 | -1.827 | 0.076 |
| Subspecies Molossinus | -0.142 | 0.021 | -6.743 | 0.000 |
| Strain G | -0.063 | 0.018 | -3.611 | 0.001 |
| Strain LEW | -0.040 | 0.020 | -2.034 | 0.050 |
| Strain PWD | -0.024 | 0.023 | -1.033 | 0.309 |
| Strain MSM | 0.140 | 0.027 | 5.169 | 0.000 |
| Strain SKIVE | -0.012 | 0.022 | -0.541 | 0.592 |
| Strain KAZ | -0.012 | 0.026 | -0.453 | 0.653 |

**Supplemental Table 30.**
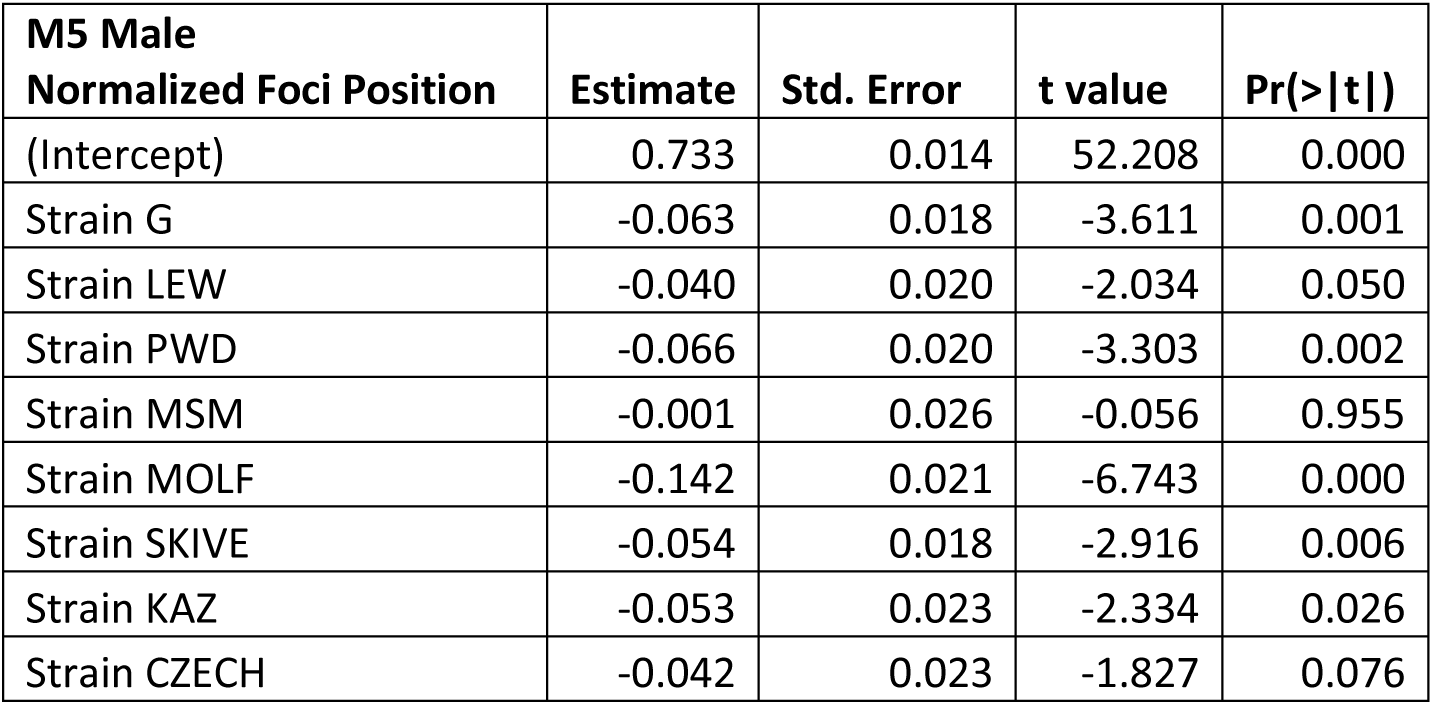

| **M5 Male**<br>**Normalized Foci Position** | **Estimate** | **Std. Error** | **t value** | **Pr(>\|t\|)** |
| --- | --- | --- | --- | --- |
| (Intercept) | 0.733 | 0.014 | 52.208 | 0.000 |
| Strain G | -0.063 | 0.018 | -3.611 | 0.001 |
| Strain LEW | -0.040 | 0.020 | -2.034 | 0.050 |
| Strain PWD | -0.066 | 0.020 | -3.303 | 0.002 |
| Strain MSM | -0.001 | 0.026 | -0.056 | 0.955 |
| Strain MOLF | -0.142 | 0.021 | -6.743 | 0.000 |
| Strain SKIVE | -0.054 | 0.018 | -2.916 | 0.006 |
| Strain KAZ | -0.053 | 0.023 | -2.334 | 0.026 |
| Strain CZECH | -0.042 | 0.023 | -1.827 | 0.076 |

**Supplemental Table 31.**

| **M1 Normalized Interfocal Distance** | **p values** |  | **Coefficients (fixed estimates)** | **Random Effects (standard deviation)** |
| --- | --- | --- | --- | --- |
| Subspecies | 0.031 | Intercept | 0.475 |  |
| Sex | 0.000 | Subspecies Musculus | 0.006 |  |
| Subspecies*Sex | 0.047 | Subspecies Molossinus | -0.003 |  |
| Strain(random) | 0.244 | Sex(male) | 0.069 |  |
|  |  | Musculus*male | 0.052 |  |
|  |  | Molossinus*male | -0.008 |  |
|  |  | Intercept (strain) |  | 0.009 |
|  |  | Strain (residual) |  | 0.048 |

**Supplemental Table 32.**

| **M2 Normalized Interfocal Distance** | **Estimate** | **Std. Error** | **t value** | **Pr(>\|t\|)** |
| --- | --- | --- | --- | --- |
| (Intercept) | 0.461 | 0.023 | 19.812 | 0.000 |
| Subspecies Musculus | 0.021 | 0.031 | 0.659 | 0.512 |
| Subspecies Molossinus | 0.003 | 0.048 | 0.061 | 0.952 |
| Sex (male) | 0.082 | 0.031 | 2.619 | 0.011 |
| Strain G | 0.027 | 0.030 | 0.892 | 0.375 |
| Strain LEW | 0.012 | 0.028 | 0.432 | 0.667 |
| Strain PWD | -0.002 | 0.026 | -0.073 | 0.942 |
| Strain MSM | 0.009 | 0.036 | 0.247 | 0.806 |
| Strain SKIVE | 0.000 | 0.031 | 0.010 | 0.992 |
| Subspecies Musculus * Sex (male) | -0.043 | 0.046 | -0.934 | 0.354 |
| Subspecies Molossinus * Sex (male) | -0.016 | 0.047 | -0.338 | 0.736 |
| Strain G * Sex (male) | -0.029 | 0.040 | -0.739 | 0.462 |
| Strain LEW * Sex (male) | 0.000 | 0.041 | -0.007 | 0.995 |
| Strain PWD * Sex (male) | 0.085 | 0.043 | 1.993 | 0.050 |
| Strain SKIVE * Sex (male) | 0.121 | 0.045 | 2.699 | 0.009 |

**Supplemental Table 33.**

| **M3 Normalized Interfocal Distance** | **Estimate** | **Std. Error** | **t value** | **Pr(>\|t\|)** |
| --- | --- | --- | --- | --- |
| (Intercept) | 0.461 | 0.023 | 19.812 | 0.000 |
| Sex (male) | 0.082 | 0.031 | 2.619 | 0.011 |
| Strain G | 0.027 | 0.030 | 0.892 | 0.375 |
| Strain LEW | 0.012 | 0.028 | 0.432 | 0.667 |
| Strain PWD | 0.019 | 0.028 | 0.668 | 0.506 |
| Strain MSM | 0.012 | 0.033 | 0.357 | 0.722 |
| Strain MOLF | -0.013 | 0.031 | -0.418 | 0.678 |
| Strain SKIVE | 0.021 | 0.033 | 0.635 | 0.528 |
| Strain KAZ | 0.021 | 0.031 | 0.659 | 0.512 |
| Strain G * Sex (male) | -0.029 | 0.040 | -0.739 | 0.462 |
| Strain LEW * Sex (male) | 0.000 | 0.041 | -0.007 | 0.995 |
| Strain PWD * Sex (male) | 0.042 | 0.041 | 1.038 | 0.303 |
| Strain MSM * Sex (male) | -0.016 | 0.047 | -0.338 | 0.736 |
| Strain SKIVE * Sex (male) | 0.078 | 0.043 | 1.821 | 0.073 |
| Strain KAZ * Sex (male) | -0.043 | 0.046 | -0.934 | 0.354 |

**Supplemental Table 34.**

| **M4 Female**<br>**Normalized Interfocal Distance** | **Estimate** | **Std. Error** | **t value** | **Pr(>\|t\|)** |
| --- | --- | --- | --- | --- |
| (Intercept) | 0.461 | 0.026 | 18.066 | 0.000 |
| Subspecies Musculus | 0.021 | 0.034 | 0.601 | 0.552 |
| Subspecies Molossinus | 0.012 | 0.036 | 0.325 | 0.747 |
| Strain G | 0.027 | 0.033 | 0.814 | 0.422 |
| Strain LEW | 0.012 | 0.031 | 0.394 | 0.696 |
| Strain PWD | -0.002 | 0.028 | -0.067 | 0.947 |
| Strain SKIVE | 0.000 | 0.034 | 0.009 | 0.993 |

**Supplemental Table 35.**

| **M5 Female**<br>**Normalized Interfocal Distance** | **Estimate** | **Std. Error** | **t value** | **Pr(>\|t\|)** |
| --- | --- | --- | --- | --- |
| (Intercept) | 0.461 | 0.026 | 18.066 | 0.000 |
| Strain G | 0.027 | 0.033 | 0.814 | 0.422 |
| Strain LEW | 0.012 | 0.031 | 0.394 | 0.696 |
| Strain PWD | 0.019 | 0.031 | 0.609 | 0.546 |
| Strain MSM | 0.012 | 0.036 | 0.325 | 0.747 |
| Strain SKIVE | 0.021 | 0.036 | 0.579 | 0.567 |
| Strain KAZ | 0.021 | 0.034 | 0.601 | 0.552 |

**Supplemental Table 36.**

| **M4 Male**<br>**Normalized Interfocal Distance** | **Estimate** | **Std. Error** | **t value** | **Pr(>\|t\|)** |
| --- | --- | --- | --- | --- |
| (Intercept) | 0.543 | 0.019 | 29.268 | 0.000 |
| Subspecies Musculus | -0.017 | 0.030 | -0.564 | 0.576 |
| Subspecies Molossinus | -0.013 | 0.028 | -0.469 | 0.642 |
| Strain G | -0.003 | 0.023 | -0.110 | 0.913 |
| Strain LEW | 0.012 | 0.026 | 0.459 | 0.649 |
| Strain PWD | 0.078 | 0.030 | 2.573 | 0.014 |
| Strain MSM | 0.009 | 0.032 | 0.277 | 0.783 |
| Strain SKIVE | 0.116 | 0.029 | 4.046 | 0.000 |
| Strain KAZ | -0.005 | 0.034 | -0.160 | 0.874 |

**Supplemental Table 37.**
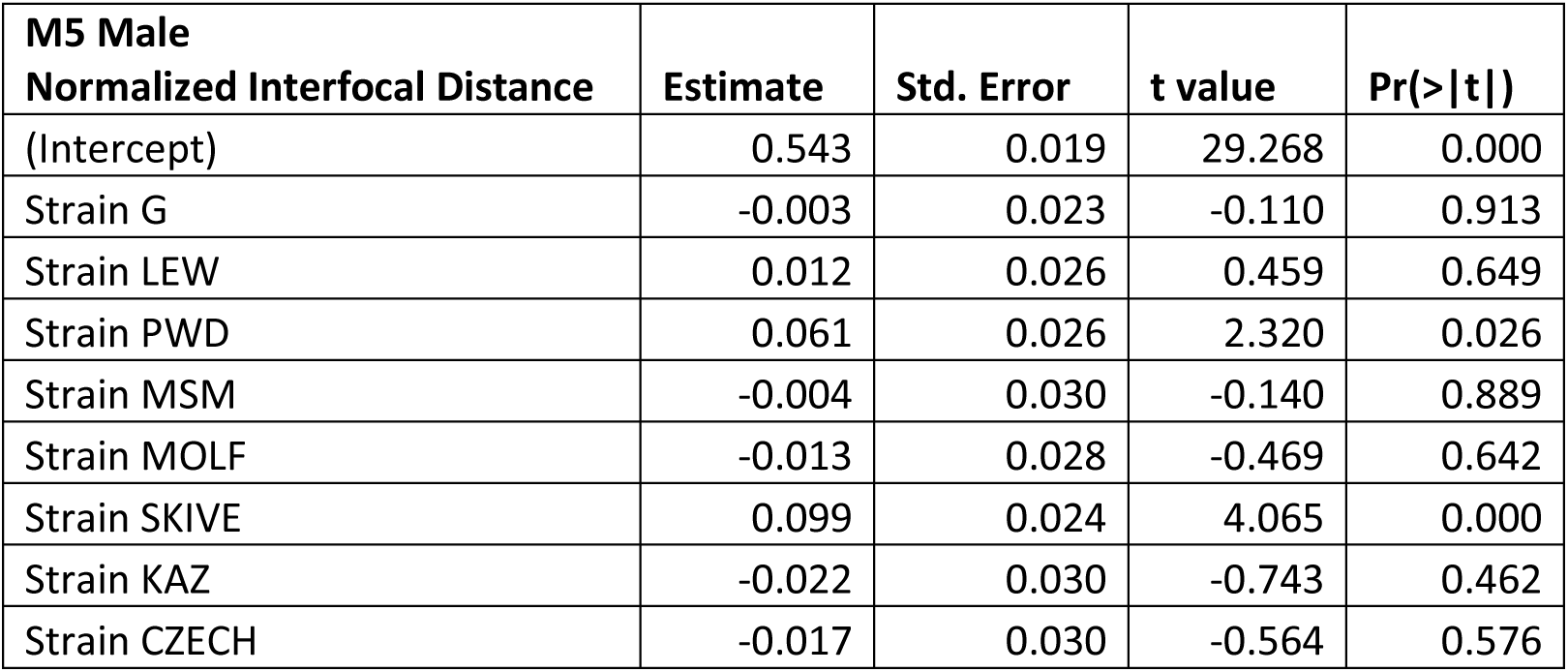

| **M5 Male**<br>**Normalized Interfocal Distance** | **Estimate** | **Std. Error** | **t value** | **Pr(>\|t\|)** |
| --- | --- | --- | --- | --- |
| (Intercept) | 0.543 | 0.019 | 29.268 | 0.000 |
| Strain G | -0.003 | 0.023 | -0.110 | 0.913 |
| Strain LEW | 0.012 | 0.026 | 0.459 | 0.649 |
| Strain PWD | 0.061 | 0.026 | 2.320 | 0.026 |
| Strain MSM | -0.004 | 0.030 | -0.140 | 0.889 |
| Strain MOLF | -0.013 | 0.028 | -0.469 | 0.642 |
| Strain SKIVE | 0.099 | 0.024 | 4.065 | 0.000 |
| Strain KAZ | -0.022 | 0.030 | -0.743 | 0.462 |
| Strain CZECH | -0.017 | 0.030 | -0.564 | 0.576 |

**Supplemental Table 38.**

| **M1 Raw**<br>**Interfocal Distance** | **p values** |  | **Coefficients**<br>**(fixed estimates)** | **Random Effects**<br>**(standard deviation)** |
| --- | --- | --- | --- | --- |
| Subspecies | 0.128 | Intercept | 57.461 |  |
| Sex | 0.026 | Subspecies Musculus | -0.713 |  |
| Subspecies*Sex | 0.084 | Subspecies<br>Molossinus | 2.932 |  |
| Strain(random) | 0.409 | Sex(male) | -6.68 |  |
|  |  | Musculus *male | 7.364 |  |
|  |  | Molossinus*male | -4.536 |  |
|  |  | Intercept (strain) |  | 0 |
|  |  | Strain (residual) |  | 7.78 |

**Supplemental Table 39.**

| **M2 Raw**<br>**Interfocal Distance** | **Estimate** | **Std. Error** | **t value** | **Pr(>\|t\|)** |
| --- | --- | --- | --- | --- |
| (Intercept) | 53.713 | 3.788 | 14.179 | 0.000 |
| Subspecies Musculus | 8.732 | 5.082 | 1.718 | 0.090 |
| Subspecies Molossinus | 4.038 | 7.886 | 0.512 | 0.610 |
| Sex (male) | -5.769 | 5.082 | -1.135 | 0.260 |
| Strain G | 6.534 | 4.890 | 1.336 | 0.186 |
| Strain LEW | 3.532 | 4.639 | 0.761 | 0.449 |
| Strain PWD | -6.918 | 4.226 | -1.637 | 0.106 |
| Strain MSM | 2.642 | 5.786 | 0.457 | 0.649 |
| Strain SKIVE | -10.073 | 5.082 | -1.982 | 0.052 |
| Subspecies Musculus * Sex (male) | -8.650 | 7.513 | -1.151 | 0.254 |
| Subspecies Molossinus * Sex (male) | -3.938 | 7.701 | -0.511 | 0.611 |
| Strain G * Sex (male) | -2.717 | 6.463 | -0.420 | 0.676 |
| Strain LEW * Sex (male) | 0.376 | 6.670 | 0.056 | 0.955 |
| Strain PWD * Sex (male) | 18.610 | 6.962 | 2.673 | 0.009 |
| Strain SKIVE * Sex (male) | 21.876 | 7.291 | 3.000 | 0.004 |

**Supplemental Table 40.**

| **M3 Raw**<br>**Interfocal Distance** | **Estimate** | **Std. Error** | **t value** | **Pr(>\|t\|)** |
| --- | --- | --- | --- | --- |
| (Intercept) | 53.713 | 3.788 | 14.179 | 0.000 |
| Sex (male) | -5.769 | 5.082 | -1.135 | 0.260 |
| Strain G | 6.534 | 4.890 | 1.336 | 0.186 |
| Strain LEW | 3.532 | 4.639 | 0.761 | 0.449 |
| Strain PWD | 1.814 | 4.553 | 0.398 | 0.692 |
| Strain MSM | 6.680 | 5.357 | 1.247 | 0.217 |
| Strain MOLF | 0.100 | 5.082 | 0.020 | 0.984 |
| Strain SKIVE | -1.341 | 5.357 | -0.250 | 0.803 |
| Strain KAZ | 8.732 | 5.082 | 1.718 | 0.090 |
| Strain G * Sex (male) | -2.717 | 6.463 | -0.420 | 0.676 |
| Strain LEW * Sex (male) | 0.376 | 6.670 | 0.056 | 0.955 |
| Strain PWD * Sex (male) | 9.960 | 6.610 | 1.507 | 0.137 |
| Strain MSM * Sex (male) | -3.938 | 7.701 | -0.511 | 0.611 |
| Strain SKIVE * Sex (male) | 13.226 | 6.956 | 1.902 | 0.062 |
| Strain KAZ * Sex (male) | -8.650 | 7.513 | -1.151 | 0.254 |

## REFERENCES

1. Anderson LK, Reeves A, Webb LM, Ashley T. 1999. Distribution of crossing over on mouse synaptonemal complexes using immunofluorescent localization of mlh1 protein. Genetics 151:1569–1579.

2. Baier B, Hunt P, Broman KW, Hassold T. 2014. Variation in genome-wide levels of meiotic recombination is established at the onset of prophase in mammalian males. PLoS genetics 10. doi:10.1371/journal.pgen.1004125

3. Bates D, Mächler M, Bolker B, Walker S. 2015. Fitting linear mixed-effects models using lme4. Journal of Statistical Software 67:1–48. doi:10.18637/jss.v067.i01

4. Baudat F, Imai Y, De Massy B. 2013. Meiotic recombination in mammals: Localization and regulation. Nature Reviews Genetics 14:794–806. doi:10.1038/nrg3573

5. Begun DJ, Aquadro CF. 1992. Levels of naturally occurring dna polymorphism correlate with recombination rates in d. Melanogaster. Nature 356:519–520. doi:10.1038/356519a0

6. Bell G. 1982. The masterpiece of nature: The evolution and genetics of sexuality. Berkeley, CA.: University of California Press.

7. Bolcun-Filas E, Schimenti J. 2012. Genetics of meiosis and recombination in mice. International review of cell and molecular biology 298:179. doi:https://doi.org/10.1016/B978-0-12-394309-5.00005-5

8. Brandvain Y, Coop G. 2012. Scrambling eggs: Meiotic drive and the evolution of female recombination rates. Genetics 190:709–723. doi:10.1534/genetics.111.136721

9. Burt A, Bell G, Harvey PH. 1991. Sex differences in recombination. Journal of evolutionary biology 4:259–277. doi:https://doi.org/10.1046/j.1420-9101.1991.4020259.x

10. Cahoon CK, Libuda DE. 2019. Leagues of their own: Sexually dimorphic features of meiotic prophase i. Chromosoma 1–16. doi:10.1007/s00412-019-00692-x

11. Charlesworth B, Morgan M, Charlesworth D. 1993. The effect of deleterious mutations on neutral molecular variation. Genetics 134:1289–1303.

12. Cutter AD, Payseur BA. 2013. Genomic signatures of selection at linked sites: Unifying the disparity among species. Nature Reviews Genetics 14:262–274. doi:https://doi.org/10.1038/nrg3425

13. Dapper AL, Payseur BA. 2017. Connecting theory and data to understand recombination rate evolution. Philosophical Transactions of the Royal Society B: Biological Sciences 372:20160469. doi:10.1098/rstb.2016.0469

14. Dumont BL, Payseur BA. 2011. Evolution of the genomic recombination rate in murid rodents. Genetics 187:643–657. doi:10.1534/genetics.110.123851

15. Dumont J, Desai A. 2012. Acentrosomal spindle assembly and chromosome segregation during oocyte meiosis. Trends in cell biology 22:241–249. doi:10.1016/j.tcb.2012.02.007

16. Felsenstein J. 1974. The evolutionary advantage of recombination. Genetics 78:737–756.

17. Fisher RA. 1930. The genetical theory of natural selection. Oxford University Press.

18. Fledel-Alon A, Leffler EM, Guan Y, Stephens M, Coop G, Przeworski M. 2011. Variation in human recombination rates and its genetic determinants. PloS one 6. doi:10.1371/journal.pone.0020321

19. Geraldes A, Basset P, Smith KL, Nachman MW. 2011. Higher differentiation among subspecies of the house mouse (mus musculus) in genomic regions with low recombination. Molecular ecology 20:4722–4736. doi:10.1111/j.1365-294X.2011.05285.x

20. Goldstein DB, Bergman A, Feldman MW. 1993. The evolution of interference: Reduction of recombination among three loci. Theoretical population biology 44:246–259. doi:10.1006/tpbi.1993.1028

21. Gruhn JR, Rubio C, Broman KW, Hunt PA, Hassold T. 2013. Cytological studies of human meiosis: Sex-specific differences in recombination originate at, or prior to, establishment of double-strand breaks. PloS one 8. doi:10.1371/journal.pone.0085075

22. Haenel Q, Laurentino TG, Roesti M, Berner D. 2018. Meta-analysis of chromosome-scale crossover rate variation in eukaryotes and its significance to evolutionary genomics. Molecular ecology 27:2477–2497. doi:10.1111/mec.14699

23. Haldane J. 1922. Sex ratio and unisexual sterility in hybrid animals. Journal of genetics 12:101–109.

24. Halldorsson BV, Palsson G, Stefansson OA, Jonsson H, Hardarson MT, Eggertsson HP, Gunnarsson B, Oddsson A, Halldorsson GH, Zink F, others. 2019. Characterizing mutagenic effects of recombination through a sequence-level genetic map. Science 363:eaau1043. doi:10.1126/science.aau1043

25. Handel MA, Schimenti JC. 2010. Genetics of mammalian meiosis: Regulation, dynamics and impact on fertility. Nature Reviews Genetics 11:124–136. doi:10.1038/nrg2723

26. Hassold T, Hunt P. 2001. To err (meiotically) is human: The genesis of human aneuploidy. Nature Reviews Genetics 2:280–291. doi:https://doi.org/10.1038/35066065

27. Hill WG, Robertson A. 1966. The effect of linkage on limits to artificial selection. Genetics Research 8:269–294. doi:https://doi.org/10.1017/S0016672300010156

28. Holloway JK, Booth J, Edelmann W, McGowan CH, Cohen PE. 2008. MUS81 generates a subset of mlh1-mlh3–independent crossovers in mammalian meiosis. PLoS genetics 4. doi:10.1371/journal.pgen.1000186

29. Hultén MA. 2011. On the origin of crossover interference: A chromosome oscillatory movement (com) model. Molecular cytogenetics 4:10. doi:10.1186/1755-8166-4-10

30. Huxley J. 1928. Sexual difference of linkage in gammarus chevreuxi. Journal of Genetics 20:145–156.

31. Inoue K, Lupski JR. 2002. Molecular mechanisms for genomic disorders. Annual review of genomics and human genetics 3:199–242. doi:10.1146/annurev.genom.3.032802.120023

32. Johnston SE, Bérénos C, Slate J, Pemberton JM. 2016. Conserved genetic architecture underlying individual recombination rate variation in a wild population of soay sheep (ovis aries). Genetics 203:583–598. doi:10.1534/genetics.115.185553

33. Koehler KE, Cherry JP, Lynn A, Hunt PA, Hassold TJ. 2002. Genetic control of mammalian meiotic recombination. I. Variation in exchange frequencies among males from inbred mouse strains. Genetics 162:297–306.

34. Kong A, Barnard J, Gudbjartsson DF, Thorleifsson G, Jonsdottir G, Sigurdardottir S, Richardsson B, Jonsdottir J, Thorgeirsson T, Frigge ML, others. 2004. Recombination rate and reproductive success in humans. Nature genetics 36:1203–1206. doi:10.1038/ng1445

35. Kong A, Thorleifsson G, Frigge ML, Masson G, Gudbjartsson DF, Villemoes R, Magnusdottir E, Olafsdottir SB, Thorsteinsdottir U, Stefansson K. 2014. Common and low-frequency variants associated with genome-wide recombination rate. Nature genetics 46:11. doi:10.1038/ng.2833

36. Kong A, Thorleifsson G, Stefansson H, Masson G, Helgason A, Gudbjartsson DF, Jonsdottir GM, Gudjonsson SA, Sverrisson S, Thorlacius T, others. 2008. Sequence variants in the rnf212 gene associate with genome-wide recombination rate. Science 319:1398–1401. doi:10.1126/science.1152422

37. Kudo NR, Anger M, Peters AH, Stemmann O, Theussl H-C, Helmhart W, Kudo H, Heyting C, Nasmyth K. 2009. Role of cleavage by separase of the rec8 kleisin subunit of cohesin during mammalian meiosis i. Journal of cell science 122:2686–2698. doi:10.1242/jcs.035287

38. Kyogoku H, Kitajima TS. 2017. Large cytoplasm is linked to the error-prone nature of oocytes. Developmental cell 41:287–298. doi:10.1016/j.devcel.2017.04.009

39. Lane S, Kauppi L. 2019. Meiotic spindle assembly checkpoint and aneuploidy in males versus females. Cellular and molecular life sciences 76:1135–1150. doi:10.1007/s00018-018-2986-6

40. Lee J. 2019. Is age-related increase of chromosome segregation errors in mammalian oocytes caused by cohesin deterioration? Reproductive Medicine and Biology. doi:10.1002/rmb2.12299

41. Lenormand T. 2003. The evolution of sex dimorphism in recombination. Genetics 163:811– 822.

42. Lenormand T, Dutheil J. 2005. Recombination difference between sexes: A role for haploid selection. PLoS biology 3. doi:10.1371/journal.pbio.0030063

43. Lenormand T, Engelstädter J, Johnston SE, Wijnker E, Haag CR. 2016. Evolutionary mysteries in meiosis. Philosophical Transactions of the Royal Society B: Biological Sciences 371:20160001. doi:10.1098/rstb.2016.0001

44. Lorch P. 2005. Sex differences in recombination and mapping adaptations. Genetica 123:39. doi:10.1007/s10709-003-2706-4

45. Lynn A, Koehler KE, Judis L, Chan ER, Cherry JP, Schwartz S, Seftel A, Hunt PA, Hassold TJ. 2002. Covariation of synaptonemal complex length and mammalian meiotic exchange rates. Science 296:2222–2225.

46. Ma L, O’Connell JR, VanRaden PM, Shen B, Padhi A, Sun C, Bickhart DM, Cole JB, Null DJ, Liu GE, others. 2015. Cattle sex-specific recombination and genetic control from a large pedigree analysis. PLoS genetics 11. doi:10.1371/journal.pgen.1005387

47. Murdoch B, Owen N, Shirley S, Crumb S, Broman KW, Hassold T. 2010. Multiple loci contribute to genome-wide recombination levels in male mice. Mammalian Genome 21:550–555. doi:https://doi.org/10.1007/s00335-010-9303-5

48. Nachman MW, Payseur BA. 2012. Recombination rate variation and speciation: Theoretical predictions and empirical results from rabbits and mice. Philosophical Transactions of the Royal Society B: Biological Sciences 367:409–421. doi:10.1098/rstb.2011.0249

49. Nagaoka SI, Hassold TJ, Hunt PA. 2012. Human aneuploidy: Mechanisms and new insights into an age-old problem. Nature Reviews Genetics 13:493–504. doi:10.1038/nrg3245

50. Otto SP, Payseur BA. 2019. Crossover interference: Shedding light on the evolution of recombination. Annual review of genetics 53:19–44. doi:10.1146/annurev-genet-040119-093957

51. Peters AH, Plug AW, Vugt MJ van, De Boer P. 1997. SHORT COMMUNICATIONS A drying-down technique for the spreading of mammalian meiocytes from the male and female germline. Chromosome research 5:66–68. doi:10.1023/A:1018445520117

52. Peterson AL, Miller ND, Payseur BA. 2019. Conservation of the genome-wide recombination rate in white-footed mice. Heredity 123:442–457. doi:10.1038/s41437-019-0252-9

53. Petkov PM, Broman KW, Szatkiewicz JP, Paigen K. 2007. Crossover interference underlies sex differences in recombination rates. Trends in Genetics 23:539–542. doi:10.1016/j.tig.2007.08.015

54. Petronczki M, Siomos MF, Nasmyth K. 2003. Un menage a quatre: The molecular biology of chromosome segregation in meiosis. Cell 112:423–440. doi:https://doi.org/10.1016/S0092-8674(03)00083-7

55. Ritz KR, Noor MA, Singh ND. 2017. Variation in recombination rate: Adaptive or not? Trends in Genetics 33:364–374. doi:10.1016/j.tig.2017.03.003

56. Samuk K, Manzano-Winkler B, Ritz KR, Noor MA. 2020. Natural selection shapes variation in genome-wide recombination rate in drosophila pseudoobscura. Current Biology. doi:10.1016/j.cub.2020.03.053

57. Sardell JM, Kirkpatrick M. 2020. Sex differences in the recombination landscape. The American Naturalist 195:361–379. doi:10.1086/704943

58. Scheipl F, Greven S, Kuechenhoff H. 2008. Size and power of tests for a zero random effect variance or polynomial regression in additive and linear mixed models. Computational Statistics & Data Analysis 52:3283–3299. doi:10.1016/j.csda.2007.10.022

59. Segura J, Ferretti L, Ramos-Onsins S, Capilla L, Farré M, Reis F, Oliver-Bonet M, Fernández-Bellón H, Garcia F, Garcia-Caldés M, others. 2013. Evolution of recombination in eutherian mammals: Insights into mechanisms that affect recombination rates and crossover interference. Proceedings of the Royal Society B: Biological Sciences 280:20131945. doi:10.1098/rspb.2013.1945

60. Shen B, Jiang J, Seroussi E, Liu GE, Ma L. 2018. Characterization of recombination features and the genetic basis in multiple cattle breeds. BMC genomics 19:304. doi:10.1186/s12864-018-4705-y

61. Smith JM, Haigh J. 1974. The hitch-hiking effect of a favourable gene. Genetics Research 23:23–35. doi:https://doi.org/10.1017/S0016672300014634

62. So C, Seres KB, Steyer AM, Mönnich E, Clift D, Pejkovska A, Möbius W, Schuh M. 2019. A liquid-like spindle domain promotes acentrosomal spindle assembly in mammalian oocytes. Science 364:eaat9557. doi:10.1126/science.aat9557

63. Subramanian VV, Hochwagen A. 2014. The meiotic checkpoint network: Step-by-step through meiotic prophase. Cold Spring Harbor perspectives in biology 6:a016675. doi:10.1101/cshperspect.a016675

64. Team R. 2015. RStudio: Integrated Development Environment for R.

65. Tease C, Hulten M. 2004. Inter-sex variation in synaptonemal complex lengths largely determine the different recombination rates in male and female germ cells. Cytogenetic and genome research 107:208–215. doi:10.1159/000080599

66. VanVeen JE, Hawley RS. 2003. Meiosis: When even two is a crowd. Current Biology 13:R831–R833. doi:10.1016/j.cub.2003.12.004

67. Wang RJ, Dumont BL, Jing P, Payseur BA. 2019. A first genetic portrait of synaptonemal complex variation. PLoS genetics 15:e1008337. doi:10.1371/journal.pgen.1008337

68. Wang RJ, Payseur BA. 2017. Genetics of genome-wide recombination rate evolution in mice from an isolated island. Genetics 206:1841–1852. doi:10.1534/genetics.117.202382

69. Wang S, Hassold T, Hunt P, White MA, Zickler D, Kleckner N, Zhang L. 2017. Inefficient crossover maturation underlies elevated aneuploidy in human female meiosis. Cell 168:977–989. doi:10.1016/j.cell.2017.02.002

70. Wong AK, Ruhe AL, Dumont BL, Robertson KR, Guerrero G, Shull SM, Ziegle JS, Millon LV, Broman KW, Payseur BA, others. 2010. A comprehensive linkage map of the dog genome. Genetics 184:595–605. doi:10.1534/genetics.109.106831

